# Optimizing the use of cryopreserved genetic resources for the selection and conservation of animal populations

**DOI:** 10.1101/2024.09.17.613440

**Authors:** Alicia Jacques, Michèle Tixier-Boichard, Gwendal Restoux

## Abstract

**Background:** Genetic diversity is essential for the sustainability and adaptability of populations, and is thus a central pillar of the agro-ecological transition. However, within a population it is inevitable that some amount of genetic variability is lost, and efforts must be made to limit this as much as possible. A valuable tool in this endeavor could be the use of cryopreserved genetic resources in cryobanks, which could assist in the management of various animal populations in the contexts of both selection and conservation.

**Results:** We performed simulations that revealed that the most appropriate use of *ex situ* genetic resources differs radically depending on characteristics of the target population and its management objectives. With populations under conservation, the aim is the management of genetic diversity, which was optimized by the regular use of cryopreserved genetic resources at each generation. For populations under selection, instead, the concern is the addition of additive genetic variability, which benefited from the use of cryopreserved collections over only a few generations based primarily on the genetic values of donors. In both cases, the use of cryopreserved individuals in animal populations requires a large amount of material: for breeds under selection because the number of offspring is high, and for breeds under conservation because the frozen semen is used repeatedly over a long period.

**Conclusions:** The use of cryopreserved material appears to be an effective means of managing the genetic variability of an animal population, either by slowing down the erosion of variability or by helping to redirect a selection objective. However, care must be taken with populations under selection to limit the disadvantages associated with the reintroduction of old genetic material, in particular the gap in breeding values for traits of interest. Finally, our study highlights the need for a sufficiently large stock of cryopreserved material in collections (e.g., number of doses, straws) to ensure the most efficient use.

## Introduction

As part of the agroecological transition aimed at improving the sustainability of animal production, efforts are currently underway to diversify farming systems at different scales (Dumont et al., 2013a). Indeed, as described by Altieri et al. (2015), the resilience of systems and their capacity to adapt are generally higher when they are characterized by mutiple levels of diversity (e.g., landscape complexity, cropping system, genetic material). Diversification has also been recommended in support of locally adapted and complementary food systems that preserve biodiversity (IPES-Food, 2016). This idea is especially relevant in the field of livestock breeding, where diversity is needed to redefine new, more sustainable breeding systems (Dumont et al., 2020; Ducos et al., 2021) that can overcome the many challenges—both environmental and societal—that have arisen in recent years (Hoffmann, 2010). For example, climate change has the potential to dramatically affect populations, thus increasing the importance of good adaptive capacities (Gaughan et al., 2019; Pasqui and Di Giuseppe, 2019). Similarly, the demand for products that are more respectful of the environment and animal welfare (i.e., local farming or free-range farming) has increased considerably in Europe, requiring a conversion of livestock production systems (Thornton, 2010; Escribano, 2016). In this context, genetic diversity and genetic resources serve as a fundamental pillar for the implementation of new breeding systems and for the maintenance of the adaptative capacities of animal populations (Notter, 1999). However, it is well known that genetic diversity tends to decrease in animal populations over time (Hagger, 2005; Eynard et al., 2016; Doekes et al., 2019; Doublet et al., 2019; Hulsegge et al., 2022). In addition to reducing the adaptive capacity of populations, a low level of genetic diversity yields slower genetic progress (Meuwissen, 1997; Sonesson et al., 2012) and an increased level of inbreeding, which the Food and Agriculture Organization recommends to limit to a 0.5%–1% increase per generation (FAO, 1998).

In order to conserve animal genetic diversity in the long term, many countries have set up gene banks to cryoconserve their genetic resources. For example, the French National Cryobank collects all of the available reproductive resources of livestock species (Danchin Burge et al., 2006). Thus far, though, existing cryopreserved resources have not been widely used in breeding programs. This may be due to a lack of knowledge, as the benefits of using cryopreserved resources in animal breeding programs have been investigated in only a few studies. Early work on this topic presented simulations showing that the use of bulls from a cryobank can help to limit the loss of genetic diversity in a population under selection, and can be used to change breeding goals (Leroy et al., 2011). Indeed, it may even be possible to use cryopreserved genetic resources to bring back alleles for adaptation to climate change, as was shown for Nordic populations (Kantanen et al., 2015). Using real data, some studies have shown the potential of using cryopreserved material to improve the short-term management of local and international breeds in the Netherlands (Doekes et al., 2018; Eynard et al., 2018). More recently, a long-term experiment using the frozen semen of an ancient bull from a French germplasm collection demonstrated the effectiveness of such a strategy in the field (Jacques et al., 2023).

We hypothesize that the limited use of cryopreserved germplasm is mainly due to a lack of available recommendations. To address this, in the present study we simulated various scenarios for the use of cryopreserved resources in different breeding schemes with different goals. In particular, we investigated the potential impact of using these resources in a local breed under conservation or for a breed under selection, with or without a change in selection objective; in each case, we identified the best strategy. This information can be used to promote the use of *ex situ* genetic resources for the management of genetic diversity.

## Methods

We performed stochastic simulations of breeding programs using the R package MoBPS, version 1.6.64 (Pook et al., 2020; Pook, 2021). All additional calculations and graphical representations were performed using R, version 4.2.1 (R Core team, 2020), and the ggplot2 R package (Wickham, 2011).

### Definition of scenarios: type of selection, prolificity and uses of germplasm collections

We defined four types of scenarios that differed in the selection process used, and considered two traits of interest, hereafter named Trait 1 and Trait 2. Both traits have a heritability of 0.4 and exhibit a negative genetic correlation of –0.3.

The scenario pop_rm corresponds to a random choice of parents among the candidates; the scenario pop_max_BV corresponds to a choice of parents that maximizes genetic progress on a synthetic index combining both traits; the scenario pop_max_DG corresponds to a choice of parents that minimizes the loss of genetic diversity; and the scenario pop_OCS corresponds to a choice of parents that maximizes genetic progress under the constraint of a maximal increase in kinship of 0.5% per generation, as recommended by the FAO (FAO, 1998). The pop_max_BV, pop_max_DG, and pop_OCS scenarios were modeled using the optimal contribution strategy (OCS) (Meuwissen, 1997) implemented in MoBPS via the optiSel package, version 2.0.5 (Wellmann, 2019; Wellmann, 2021). The relationship matrix in the OCS was calculated using the vanRaden method (VanRaden, 2008). For the scenarios involving the use of OCS, we set a uniform constraint for the female kernel, allowing us to consider their contributions to be equal. The maximum contribution of each male candidate was set at 5% for the pop_max_BV scenario, and 1% for the pop_max_DG and pop_OCS scenarios, in order to obtain a realistic minimum number of fathers at each generation. For the pop_OCS scenario, an upper bound to restrict the average kinship of the progeny was calculated for each generation, based on an estimate of average kinship within the male and female kernels proposed as candidates and prohibiting an increase beyond 0.5%.

The effect of prolificity was studied by simulating three levels: 1, 2, or 10 offspring per mating. These values were chosen to cover the majority of livestock species (ruminants, horses, and pigs).

Finally, these scenarios were implemented either with the contemporary kernels only, or by adding 40 cryopreserved sires to the pool of candidates. In the latter case, cryopreserved individuals were systematically re-evaluated at the same time as contemporary individuals.

### Definition of scenarios: uses of germplasm collections during a change in breeding goal

This case applied only to populations under selection. Three categories of breeding goal changes were defined, which differed in the weights given to Trait 1 and Trait 2 in the synthetic index. The first category corresponded to a constant selection objective in which Trait 1 had the highest weight (0.8); the second category corresponded to a moderate change in selection objective, in which each trait had an equal weight (0.5); and the last category corresponded to a strong change in objective, with a complete reversal of the weights of Trait 1 and Trait 2 in the synthetic index (i.e.,*weight*_*Trait*1_ = 0.2 and *weight*_*Trait2*_ = 0.8). Each category of breeding goal change was tested for the three levels of prolificity. We simulated 15 generations following pre-burn-in and the selection burn-in (i.e., beginning with the creation of the 21st generation).

### Initialization of simulated populations and pre-burn-in

#### Genetic architecture and definition of a synthetic index

First, we simulated a founder population of 500 individuals with a sex ratio of 0.5, with 5000 genetic markers (SNPs) distributed over 4 chromosomes. SNPs were regularly distributed along the genome, i.e., the average distance in centimorgans between neighboring SNPs was 4.105 cM.

Each trait was simulated with 500 additive quantitative trait loci (QTLs) that were randomly distributed along the genome, the effects of which were drawn from a Gaussian distribution. The parameters for the two traits were initialized with a genetic mean of 0 and a variance of 1. The true breeding values (BVs) were generated by the genotype simulation and used to standardize the EBV for each trait. Then, EBVs were combined in a synthetic index using a weight of 0.8 for Trait 1 and a weight of 0.2 for Trait 2. Thus, for each animal, the index value was obtained as follows:

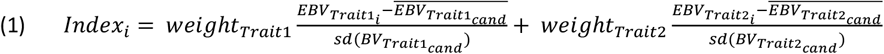

with *Index*_*i*_ the synthetic index value of animal *i*; 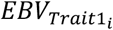 and 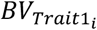 the estimated breeding value and the true breeding value, respectively, of animal *i* for Trait 1 (idem for Trait 2); and *weight*_*Trait1*_and *weight*_*Trait*2_ the weights of Trait 1 and Trait 2 in the selection objective, respectively (here, 0.8 and 0.2).

The candidate sires were ranked according to this index. The estimated genetic values of individuals (EBVs) were obtained using GBLUP evaluation models. The SNPs involved in the QTLs of the two traits were not taken into account in the evaluations, which is generally the case with the SNP chips used routinely in breeding schemes as the causal SNPs are not available. The reference population for the estimates consisted of all phenotyped individuals available at the time of evaluation.

#### Pre-burn-in and initial population creation

We performed a pre-burn-in process to generate linkage disequilibrium and genetic structure for 20 simulated populations (corresponding to the 20 replicates for the different scenarios that follow). To do this, we simulated 20 founder kernels of 250 males and 250 females, then random breeding cycles for 10 generations. Each new cohort included 250 males and 250 females (candidates for selection) resulting from the mating of the 20 best males and the 250 best females of the previous cohort, with prolificity set at 2, half-sib matings prohibited, and the selection of sires based on the synthetic index defined above (Figure 1). At the end of these 10 generations, the aim was to obtain a population with an average kinship of 10% in generation 10.

**Figure 1.**
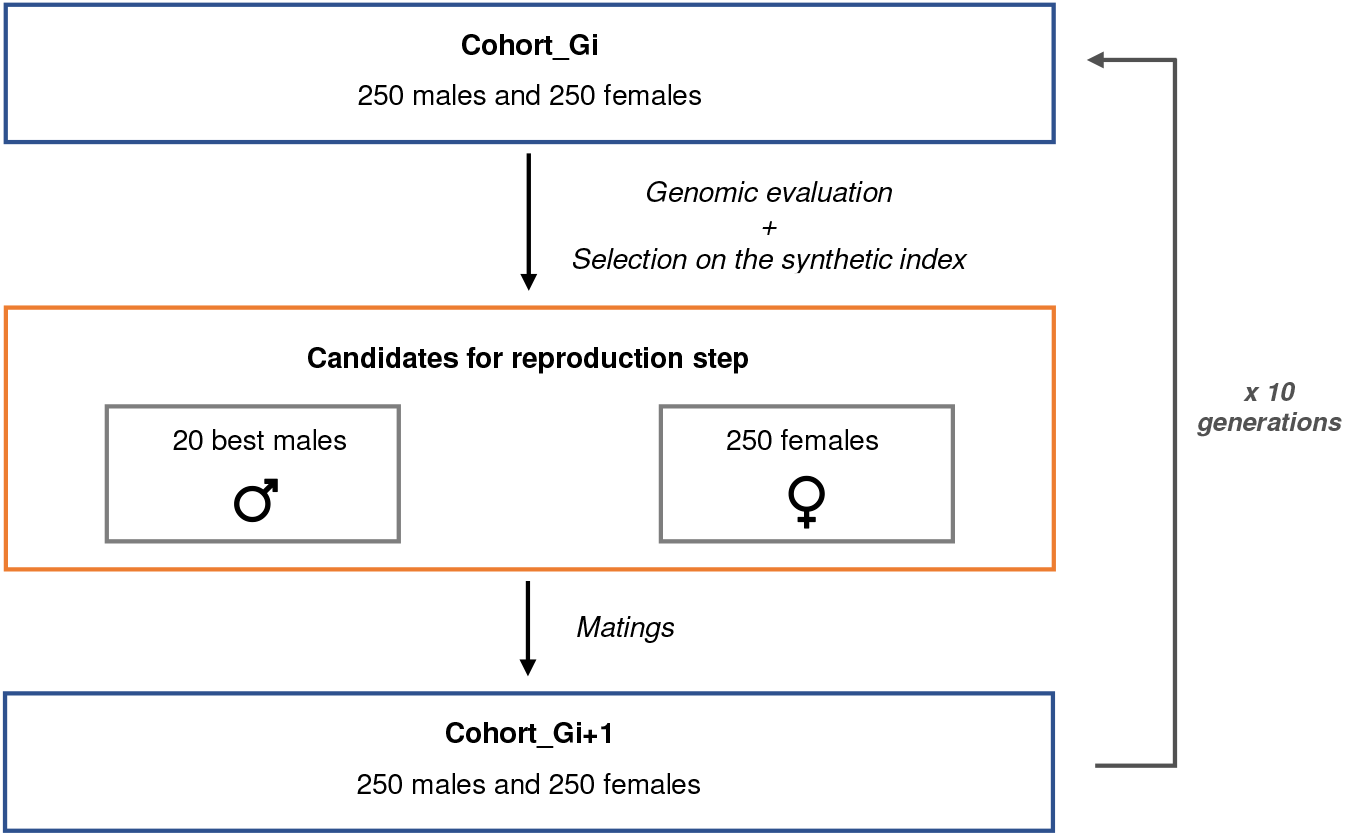
Breeding steps for the pre-burn-in phase of simulations.

**Figure 2.**
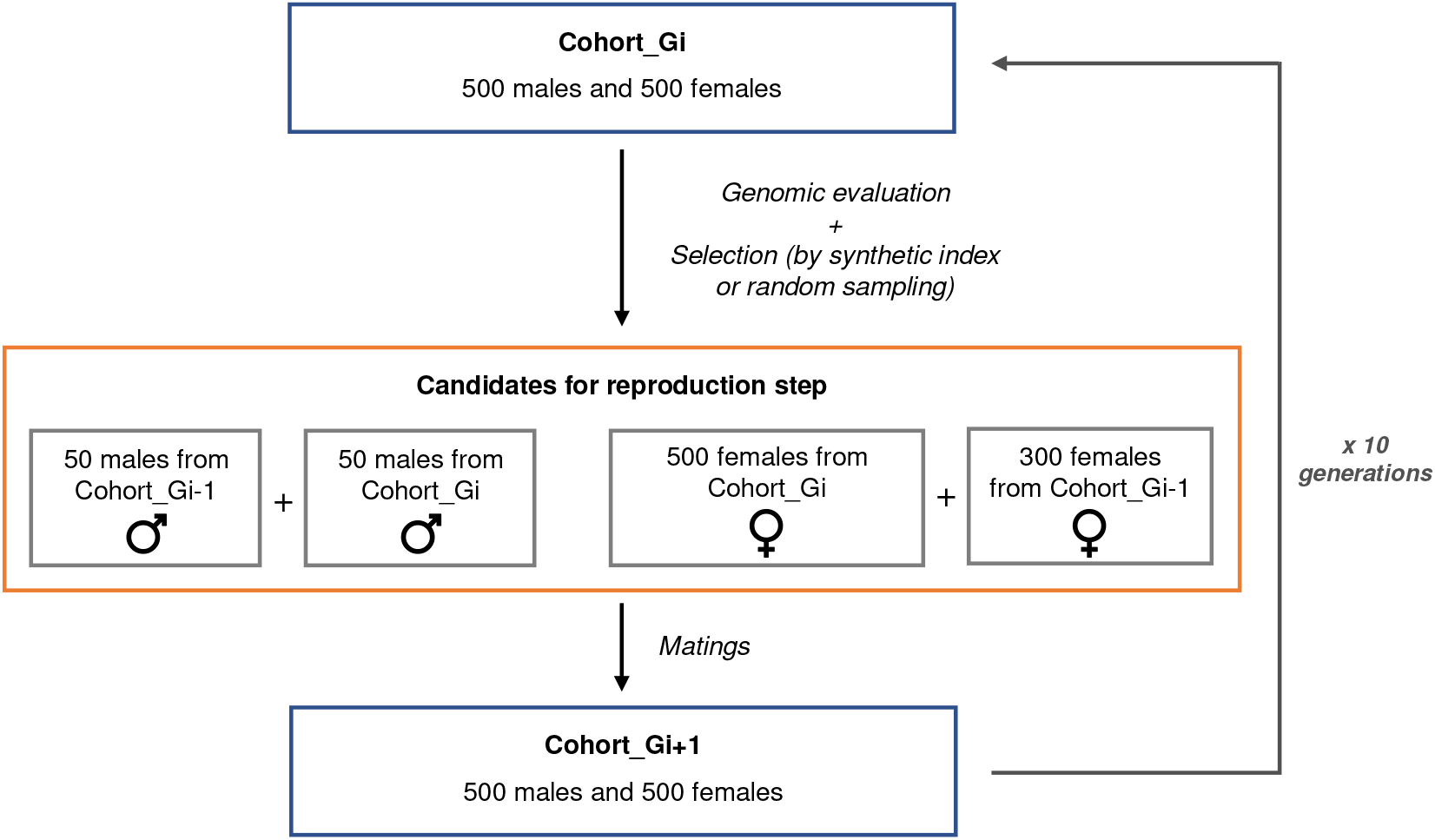
Breeding steps for the burn-in phase for populations under selection or conservation.

### Burn-in phases, breeding scheme, and definition of germplasm collection

#### Burn-in and simulated population size

Following the pre-burn-in phase, we performed two types of burn-in: one to mimic a population under selection and the other to mimic a population under conservation.

We simulated a breeding program for livestock with overlapping generations. Each new cohort was composed of 500 males and 500 females, obtained from matings between a kernel of 100 males and 800 females from the last two generations, with prolificity always set at 2. Individuals were chosen for reproduction (i) on the basis of the synthetic index defined above for selection burn-in or (ii) by random sampling in the case of conservation burn-in (in which case the only influence on genetic diversity was genetic drift). These burn-in processes were performed over 10 generations to obtain an average kinship in the 20^th^ generation of (i) 14–15% in the case of the selection scheme or (ii) 10–11% in the case of the conservation scheme.

#### Simulation of cryopreserved individuals: creation of germplasm collection

From the 50 individuals proposed as potential sires for each generation, 10 were chosen from cryopreserved samples. Thus, cryopreserved individuals may have already been used as sires before, which is generally the case for the sires present in the French National Cryobank. The simulated cryobank contained 10 cryopreserved sires from generations 11, 13, 15, and 17; these 40 individuals represented a rough approximation of the current heterogeneity of the French National Cryobank (Jacques et al., 2024). A summary of the different simulation steps is presented in Additional File 1 Figure S1.

### Indicators of scenarios, genetic diversity, and uses of germplasm collections

#### Quantification of the use of cryopreserved individuals

For each scenario involving the use of *ex situ* genetic resources, we quantified the effective use of cryopreserved individuals by counting their average number of direct descendants (i.e., sons or daughters), as well as indirect descendants (e.g., grandchildren, great-grandchildren). Thus, for each rank (the number of generations separating a cryopreserved father from his descendants, e.g., rank 1 for children, rank 2 for grandchildren, rank 3 for great-grandchildren, etc.), the number of descendants resulting from the use of cryopreserved individuals was recorded.

#### Measurements of performance

For each generation, the “true” genetic values (BVs) of the two traits were calculated, together with the empirical genetic variances for these two traits (i.e., variance of the genetic values of the simulated individuals). The EBVs for traits 1 and 2, as well as the value of the sires’ standardized synthetic index, were evaluated at each generation. We also calculated a value for the “true” selection index, i.e., based on EBVs and without standardization, for each generation by applying their respective weights in the selection objective. In this way, we were able to compare the simultaneous evolution of the two traits.

#### Measurements of genetic diversity

The evolution of genetic diversity was studied by comparing its value at the start of the program (generation 20) with that obtained at the end of the program (generation 35). To do this, we extracted the genotypes of 1,000 individuals from generations 20 and 35 for each simulation, and estimated various measures of diversity using PLINK 1.9 software (Purcell et al., 2007; Chang et al., 2015).

First, we measured the mean heterozygosity values at these two time steps (HeG20 and HeG35), to obtain the following differential:

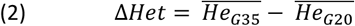

In a similar way, we investigated variations in allele frequencies over time between generation 20 and generation 35 of each simulation using PLINK’s “freq” function. We then calculated the mean difference between the allelic frequencies in generation 35 and generation 20 for all SNPs according to the following equation:

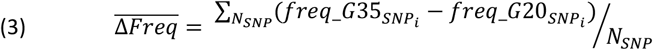

with 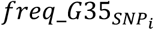 the frequency of marker *i* in the 35th generation, 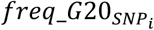 the frequency of marker *i* in the 20th generation, and *N*_*SNP*_the total number of markers considered.

We also examined the number of enriched rare SNPs, defined as those with a minor allele frequency (MAF) below 5% in the G20 cohort and above 5% in the G35 cohort.

Average kinship was calculated for each generation using the “kinship.emp.fast” function of the MoBPS package, which estimates the real kinship based on the recombination points of genotypes by sampling the population. This estimator is based on the notion of shared identical segments per progeny. From this, the increase in average kinship (Δ*K*) was computed.

Finally, the genetic distance between generation 20 and generation 35 was estimated for each scenario using Nei’s method (Nei, 1972) with MoBPS’s “get.distance” function.

### Statistical analyses

Statistical tests were performed using the “lm” function, post-hoc tests were conducted using the emmeans package (Lenth et al., 2021), and type II ANOVAs were performed using the car package (Fox et al., 2019).

## Results

### Quantifying the use of *ex situ* resources

When proposed as candidates, almost all individuals from *ex situ* collections were used in programs aimed at maintaining genetic diversity (i.e., pop_rm and pop_max_DG) (see Table 1) and between these two scenarios there was no significant difference in the number of cryobanked individuals used (two-factor ANOVA, F= 2.11, df=114, p=0.07). On the other hand, very few cryopreserved individuals were used when the objective was selection, although more were used in the pop_OCS program which took into account both diversity and selection. In this case, the number of individuals used increased with a change in objective (in proportion to the magnitude of the change; Table 1), but was not influenced by prolificacy (three-factor ANOVA, F=0.93, df=2, p=0.39). In populations under selection, cryobanked individuals were mainly used at the start of the program, whereas in populations under conservation, they were used more regularly (see Figure 3). Moreover, the number of offspring derived from cryopreserved individuals was greater in conservation populations than in selection populations (Table 1 and Figure 4). In the former, the average number of offspring per cryopreserved sire was higher in the scenario maximizing genetic diversity (i.e., pop_max_DG) (two-factor ANOVA test, F= 537.8, df=114, p<0.05). In the latter case, instead, we observed an increase in the number of direct descendants in populations for which there was no constraint on genetic diversity (pop_max_BV) compared to populations with this constraint (pop_OCS). Moreover, the number of offspring seemed to increase with the magnitude of a change in objective (from null to strong).

**Table 1.** Number of cryopreserved individuals used and direct offspring produced in populations under conservation (pop_rm and pop_max_DG) or selection (pop_max_BV and pop_OCS).

| SCENARIO TYPE | pop_rm |  | pop_max_DG |  | pop_OCS |  | pop_max_BV |  |
| --- | --- | --- | --- | --- | --- | --- | --- | --- |
|  | No. cryopreserved sires used | No. offspring per cryopreserved sire | No. cryopreserved sires used | No. offspring per cryopreserved sire | No. cryopreserved sires used | No. offspring per cryopreserved sire | No. cryopreserved sires used | No. offspring per cryopreserved sire |
| **Prolificity 1** |  |  |  |  |  |  |  |  |
| None | 40<br>[sd=0] | 107.5<br>[sd=2.24] | 40<br>[sd=0] | 162.5<br>[sd=4.25] | 0.30<br>[sd=0.47] | 23<br>[sd=10.84] | 0<br>[sd=0] | NA |
| Moderate | NA | NA | NA | NA | 1<br>[sd=0.79] | 25.86<br>[sd=14.90] | 0.20<br>[sd=0.52] | 95.5<br>[sd=2.29] |
| Strong | NA | NA | NA | NA | 2.70<br>[sd=2.13] | 31.1<br>[sd=10.42] | 0.75<br>[sd=0.78] | 138.7<br>[sd=57.96] |
| **Prolificity 2** |  |  |  |  |  |  |  |  |
| None | 40<br>[sd=0] | 107.8<br>[sd=2.41] | 40<br>[sd=0] | 160<br>[sd=5.19] | 0.25<br>[sd=0.44] | 27.2<br>[sd=10.83] | 0.05<br>[sd=0.22] | 108<br>[sd=NA] |
| Moderate | NA | NA | NA | NA | 1<br>[sd=0.97] | 28.5<br>[sd=18.73] | 0.20<br>[sd=0.52] | 113<br>[sd=29.44] |
| Strong | NA | NA | NA | NA | 2.25<br>[sd=1.48] | 27.2<br>[sd=10.94] | 0.55<br>[sd=0.83] | 136.7<br>[sd=49.91] |
| **Prolificity 10** |  |  |  |  |  |  |  |  |
| None | 40<br>[sd=0] | 109.9<br>[sd=4.23] | 39.90<br>[sd=0.31] | 152.6<br>[sd=9.81] | 0.35<br>[sd=0.59] | 25<br>[sd=18.70] | 0<br>[sd=0] | NA |
| Moderate | NA | NA | NA | NA | 0.80<br>[sd=0.89] | 31.4<br>[sd=13.43] | 0<br>[sd=0] | NA |
| Strong | NA | NA | NA | NA | 2.30<br>[sd=1.78] | 41.1<br>[sd=13.98] | 0.55<br>[sd=0.69] | 132.8<br>[sd=60.47] |

Note: Scenario abbreviations correspond to those described in the Methods section.

**Figure 3.**
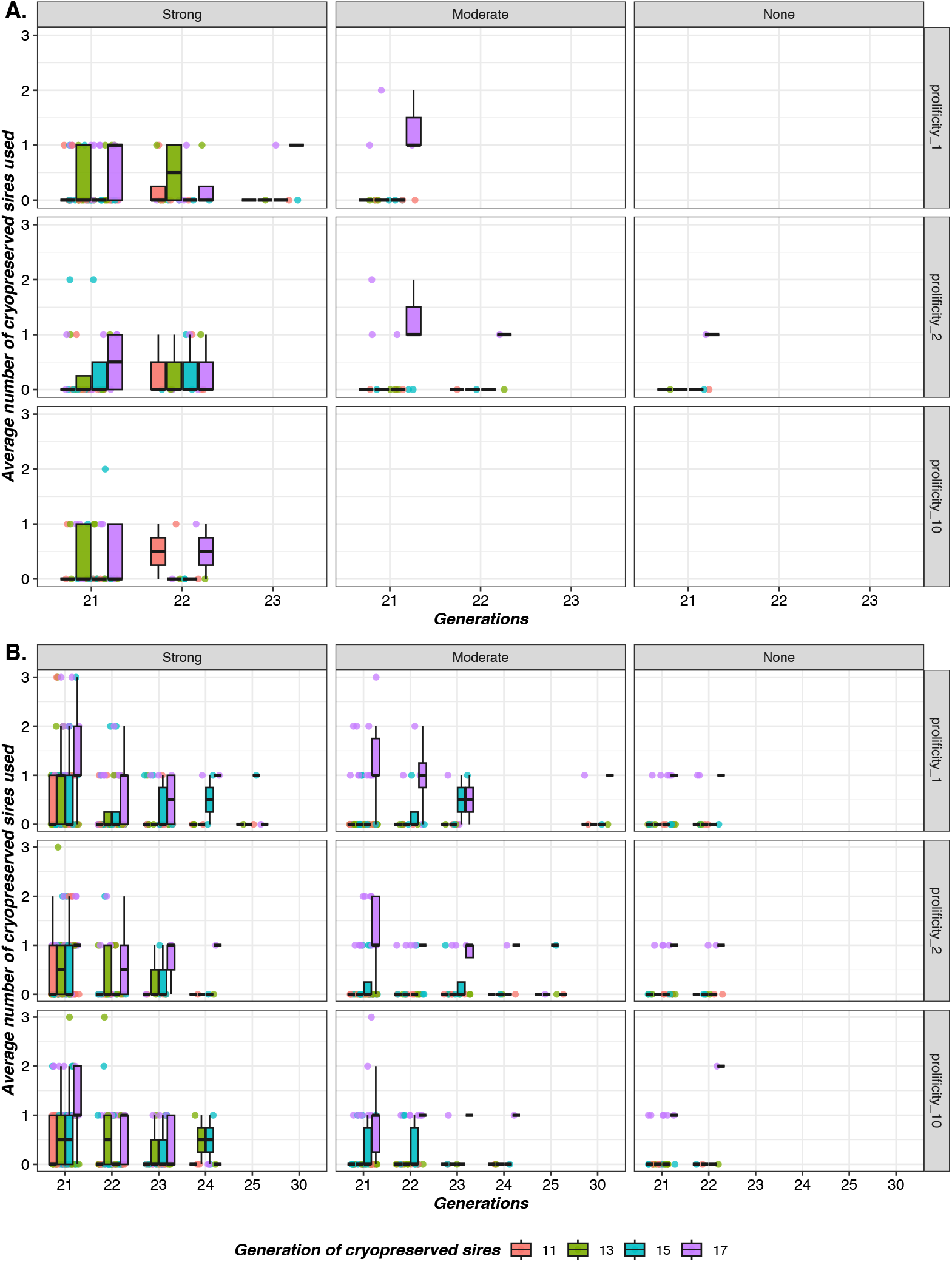
Generations in which cryopreserved genetic resources were mobilized in breeding schemes aimed at maximizing genetic progress (A) or in breeding schemes aimed at maximizing genetic progress with a constraint on genetic diversity (B). None: scenario with no change in the weighting of the two traits; moderate: weighting of both traits increased to 0.5; strong: complete inversion of weights between the two traits. In red, cryobank individuals born in generation 11; in green, cryobank individuals born in generation 13; in blue, cryobank individuals born in generation 15; in violet, cryobank individuals born in generation 17.

**Figure 4.**
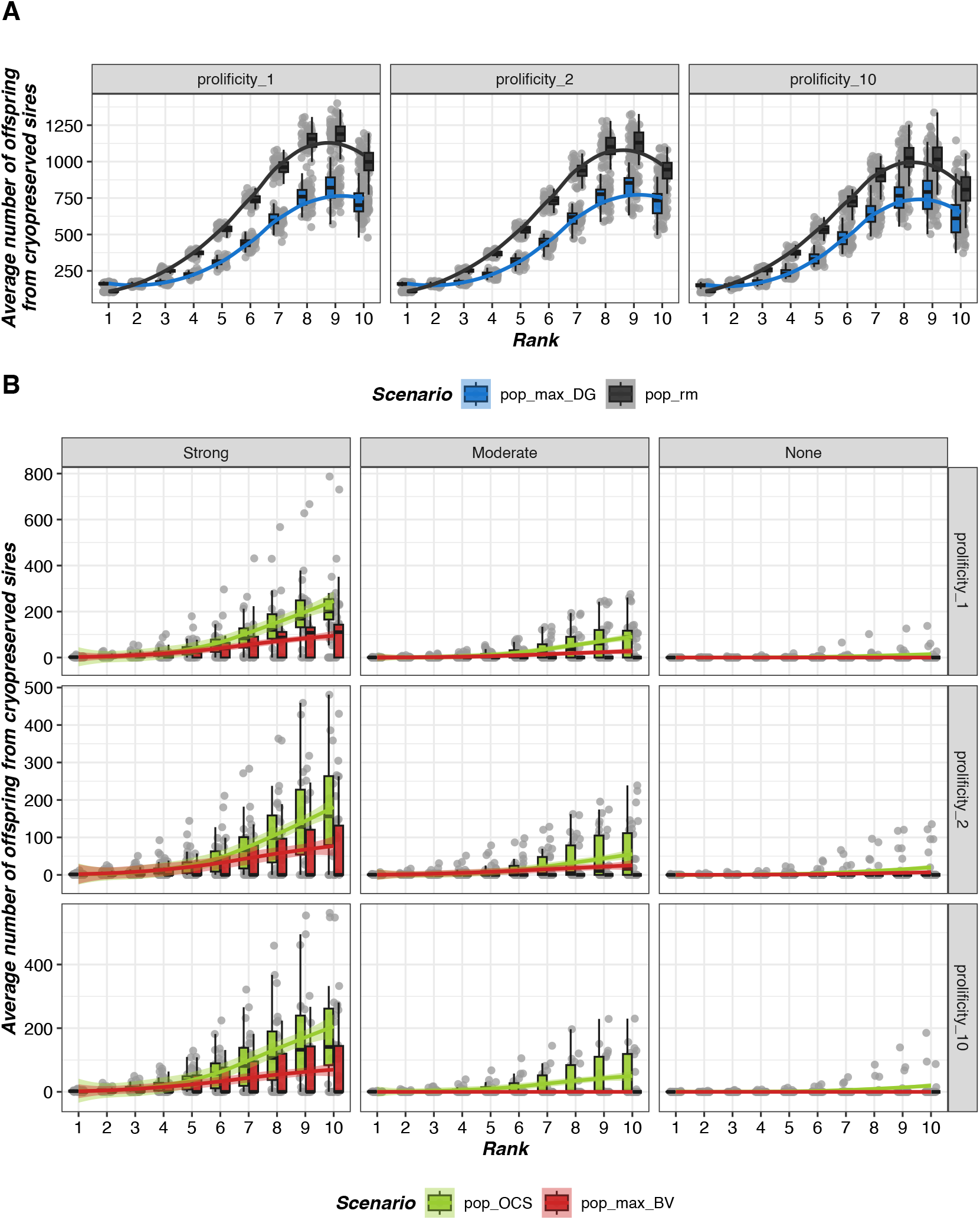
Number of offspring of cryopreserved individuals in the different management programs and scenarios across generations. The x-axis corresponds to the genealogical rank of individuals in relation to their ancestors from the *ex situ* collections (i.e., rank 1 for children, rank 2 for grandchildren, rank 3 for great-grandchildren, etc.).

In the long term, the overall contribution of individuals from cryopreserved collections was significant, with family origins persisting over generations, especially when selection pressure was relaxed (see Figure 4).

### Impact on genetic diversity

In most cases, the average kinship in a population increased less rapidly when cryopreserved collections were used, particularly when the objective was only genetic diversity (i.e., pop_rm, pop_max_DG) (three-factor ANOVA, F=622.3, df=230, p<0.05) or genetic progress (i.e., pop_max_BV) (three-factor ANOVA, F=521.25, df=1, p<0.05). Instead, no difference was noted in average kinship in the pop_OCS program (see Figure 5 and Figure 8). The same pattern was observed for the average observed heterozygosity of the populations, which, although inevitably decreasing over time, was less degraded in the scenarios using *ex situ* collections (three-factor ANOVA, F=465.20, df=1, p<0.05 for pop_max_BV and F=242, df=230, p<0.05 for pop_rm and pop_max_DG). Again, though, the exception was pop_OCS, for which the difference was only visible when prolificity was set to 10 (Figure 8). Similar findings were noted in the evolution of allelic frequencies and MAF, which declined less when using cryopreserved collections (i.e., Mean_Delta_Freq) (three-factor ANOVA, F=442.47, df=1, p<0.05 for pop_max_BV and F=228.3, df=230, p<0.05 for conservation scenarios), resulting in a smaller decrease in expected heterozygosity (i.e., He=2×MAF×(1-MAF)). The use of cryopreserved genetic resources led to a significant increase in the number of rare SNPs conserved in populations (three-factor ANOVA test, F=134.31, df=1, p-<0.05 for pop_max_BV and F= 100.4, df=230, p<0.05 for pop_rm and pop_max_DG). Details of the measurements of genetic diversity for the different scenarios are presented in Additional File 2 (Table S1, Table S2 for populations under selection, and Table S3 for populations under conservation).

**Figure 5.**
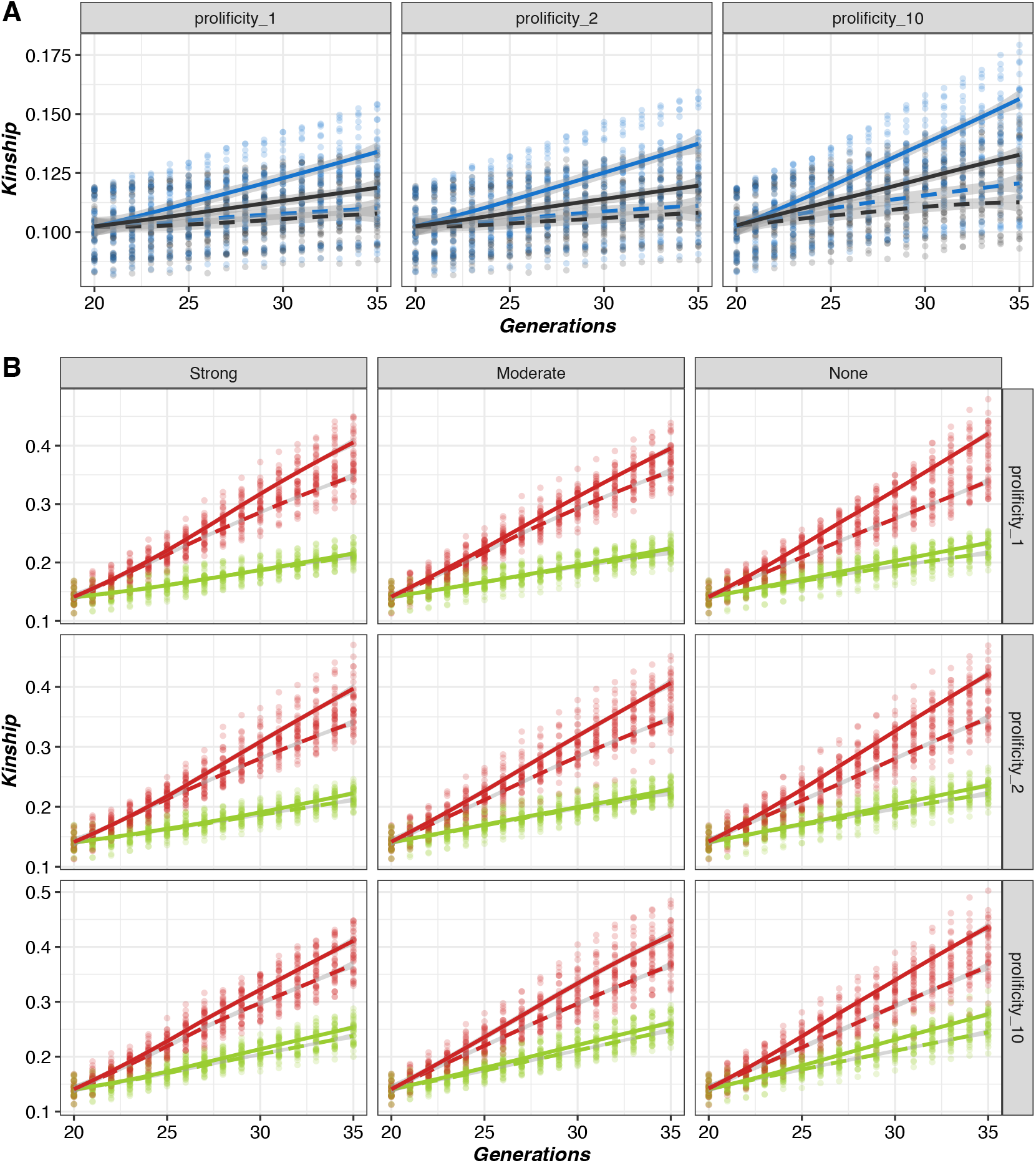
Evolution of average kinship across generations within populations in the different scenarios, with a diversity objective only (A) or with a selection objective (B). The solid and dashed lines represent results without and with the use of *ex situ* collections, respectively.

Populations differentiated less over time when individuals from *ex situ* collections were used, with lower Nei distances between generations 20 and 35 in all scenarios (see Figure 6, 7, and 8) (three-factor ANOVA, F=525.15, df=1, p<0.05 for pop_max_BV and F=982, df=230, p<0.05 for conservation scenarios).

**Figure 6.**
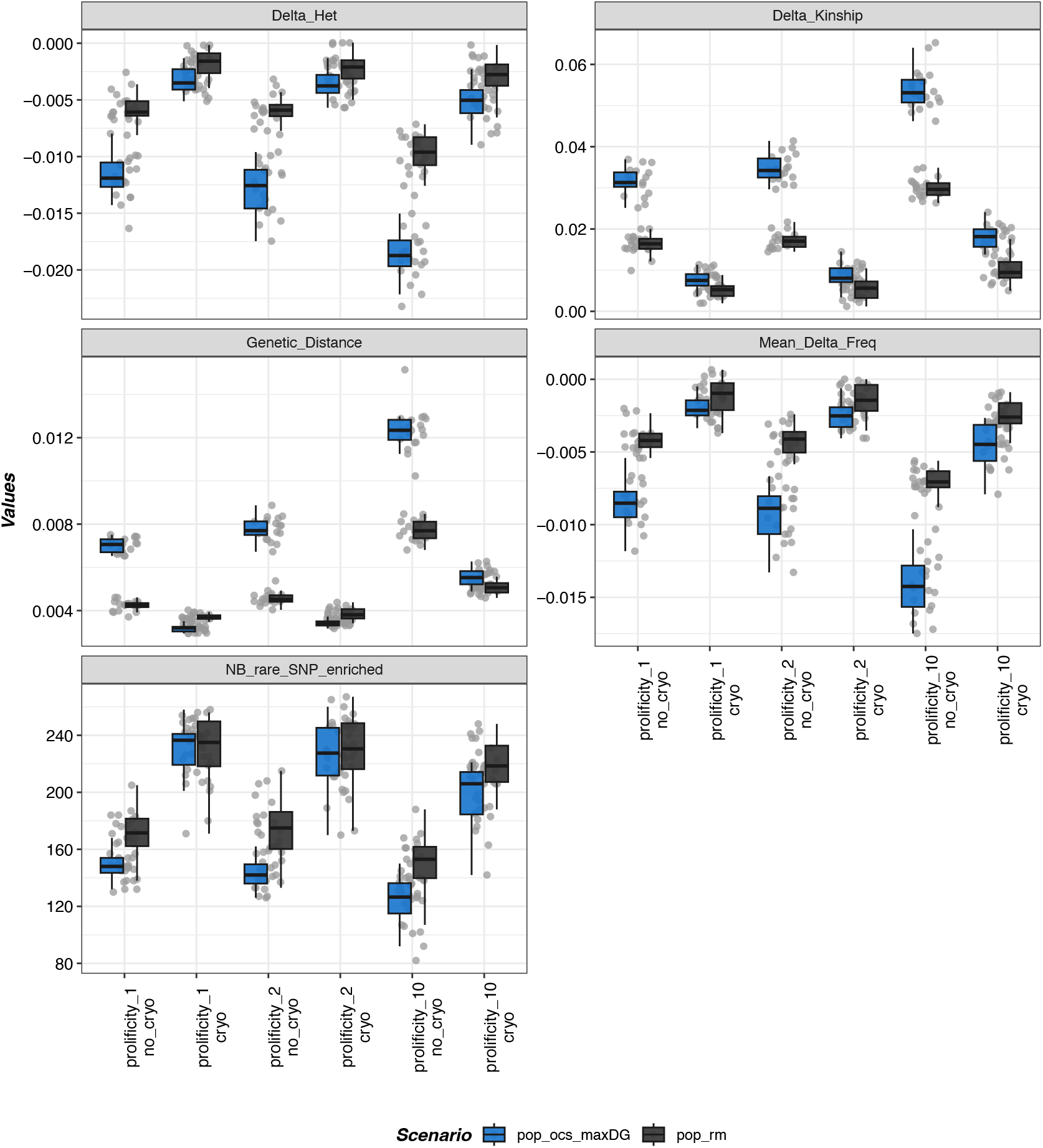
Measurements of genetic diversity for scenarios of conservation populations. In black, scenario with random mating (pop_rm). In blue, scenario maximizing genetic diversity (pop_max_DG). Each dot represents one replicate. Delta_Het: the difference in heterozygosity between generations 20 and 35; Delta_Kinship: the difference in kinship between generations 20 and 35; Genetic_Distance: Nei’s genetic distance between generations 20 and 35; Mean_Delta_Freq: the difference in allelic frequencies between generations 20 and 35; NB_rare_SNP_enriched: the number of SNPs with a MAF lower than 0.05 in generation 20 whose frequency increased in generation 35.

**Figure 7.**
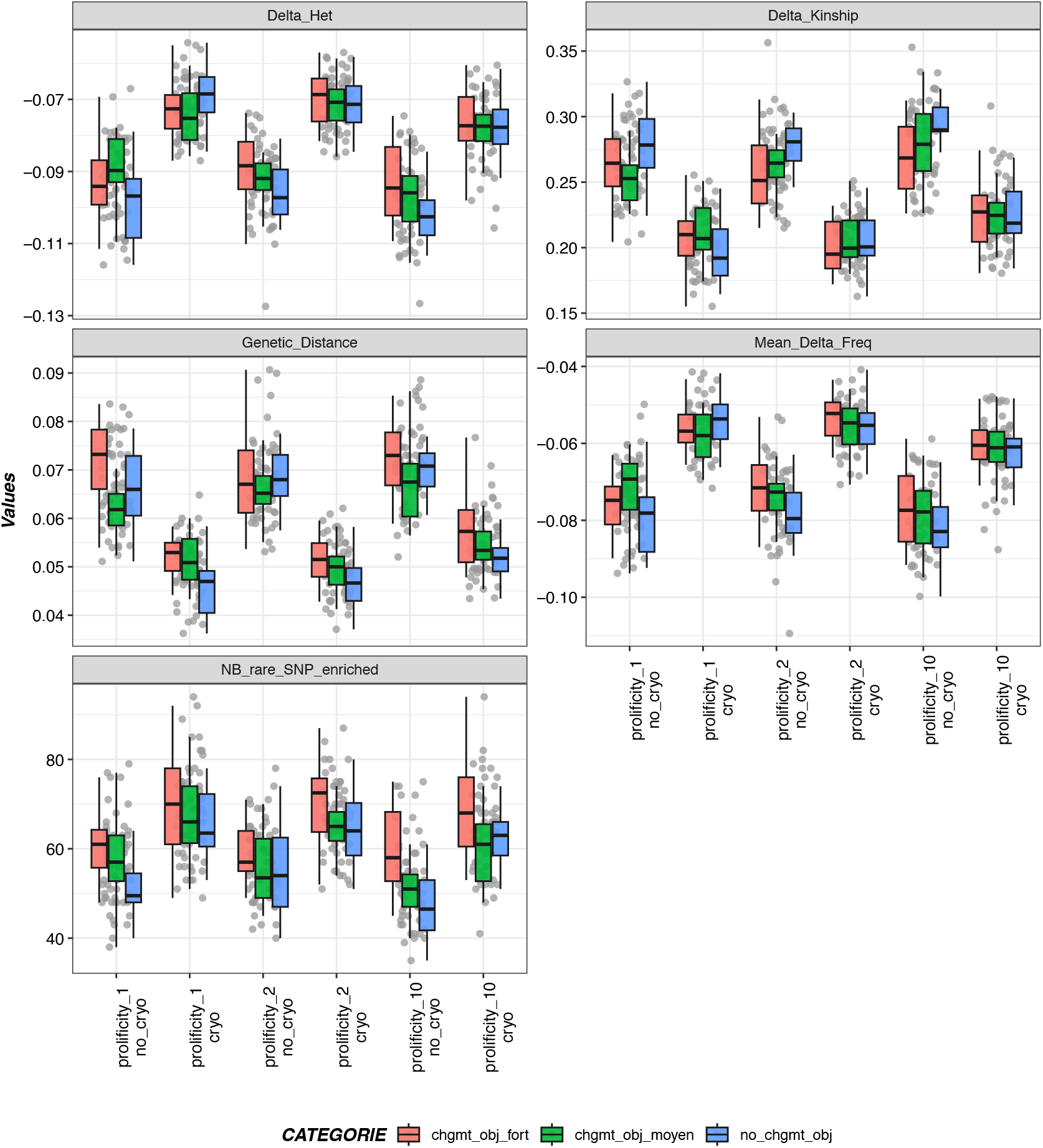
Measurements of genetic diversity for pop_max_BV scenarios. In blue, none: scenario with no change in the weighting of the two traits; in green, moderate: weighting of both traits increased to 0.5; in red, strong: complete inversion of weights between the two traits. Delta_Het: the difference in heterozygosity between generations 20 and 35; Delta_Kinship: the difference in kinship between generations 20 and 35; Genetic_Distance: Nei’s genetic distance between generations 20 and 35; Mean_Delta_Freq: the difference in allelic frequencies between generations 20 and 35; NB_rare_SNP_enriched: the number of SNPs with a MAF lower than 0.05 in generation 20 whose frequency increased in generation 35.

**Figure 8.**
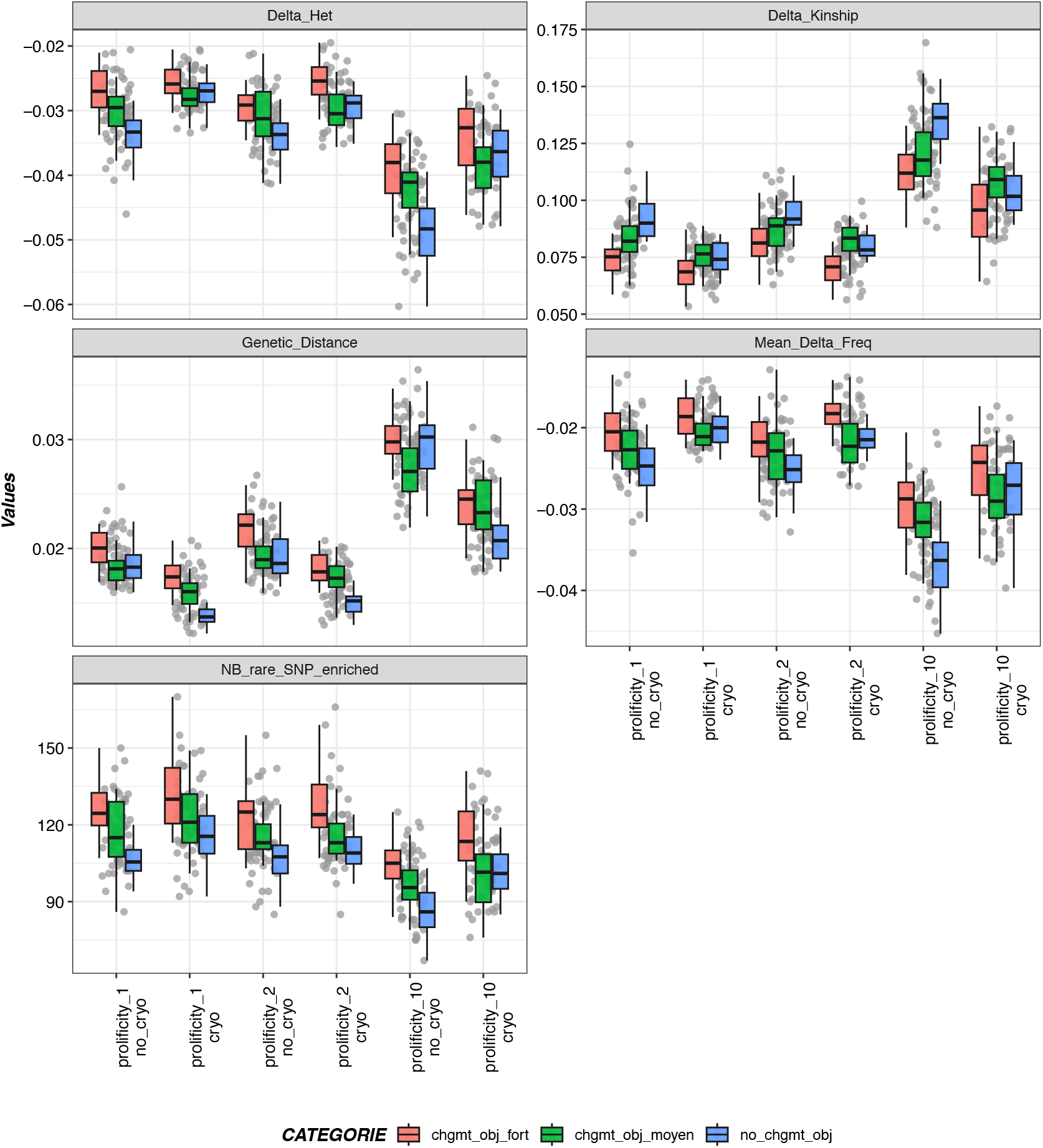
Measurements of genetic diversity for pop_OCS scenarios. In blue, none: scenario with no change in the weighting of the two traits; in green, moderate: weighting of both traits increased to 0.5; in red, strong: complete inversion of weights between the two traits. Delta_Het: the difference in heterozygosity between generations 20 and 35; Delta_Kinship: the difference in kinship between generations 20 and 35; Genetic_Distance: Nei’s genetic distance between generations 20 and 35; Mean_Delta_Freq: the difference in allelic frequencies between generations 20 and 35; NB_rare_SNP_enriched: the number of SNPs with a MAF lower than 0.05 in generation 20 whose frequency increased in generation 35.

### Impact on genetic values

The results presented in this section focus only on scenarios involving genetic progress in their objectives (i.e., pop_max_BV and pop_OCS). Overall, the impact of using *ex situ* collections on the value of the synthetic index was not significant (three-factor ANOVA, F=1.24, df=2, p=0.29 for pop_OCS and F=0.62, df=2, p=0.54 for pop_max_BV) (see Figure 9). When we looked at the two traits comprising the index, though, we observed differences in behavior based on changes in the selection objective (Figure 10). There was no effect when the change of objective was moderate. Instead, in cases in which there was no change in objective, the use of cryopreserved collections enabled trait 2 to be maintained or even improved (in contrast to the deterioration that occured without *ex situ* contributions), while trait 1 deteriorated in pop_max_BV programs and was relatively unchanged in pop_OCS programs. In cases in which the change in objective was significant, we observed the opposite pattern: a positive impact of *ex situ* collections on trait 1, together with a slight degradation of trait 2 in pop_max_BV scenarios and no impact in pop_OCS programs (four-factor ANOVA, F=0.25, df=2, p=0.76 for Trait 1 and F=1.05, df=2, p=0.35 for Trait 2).

**Figure 9.**
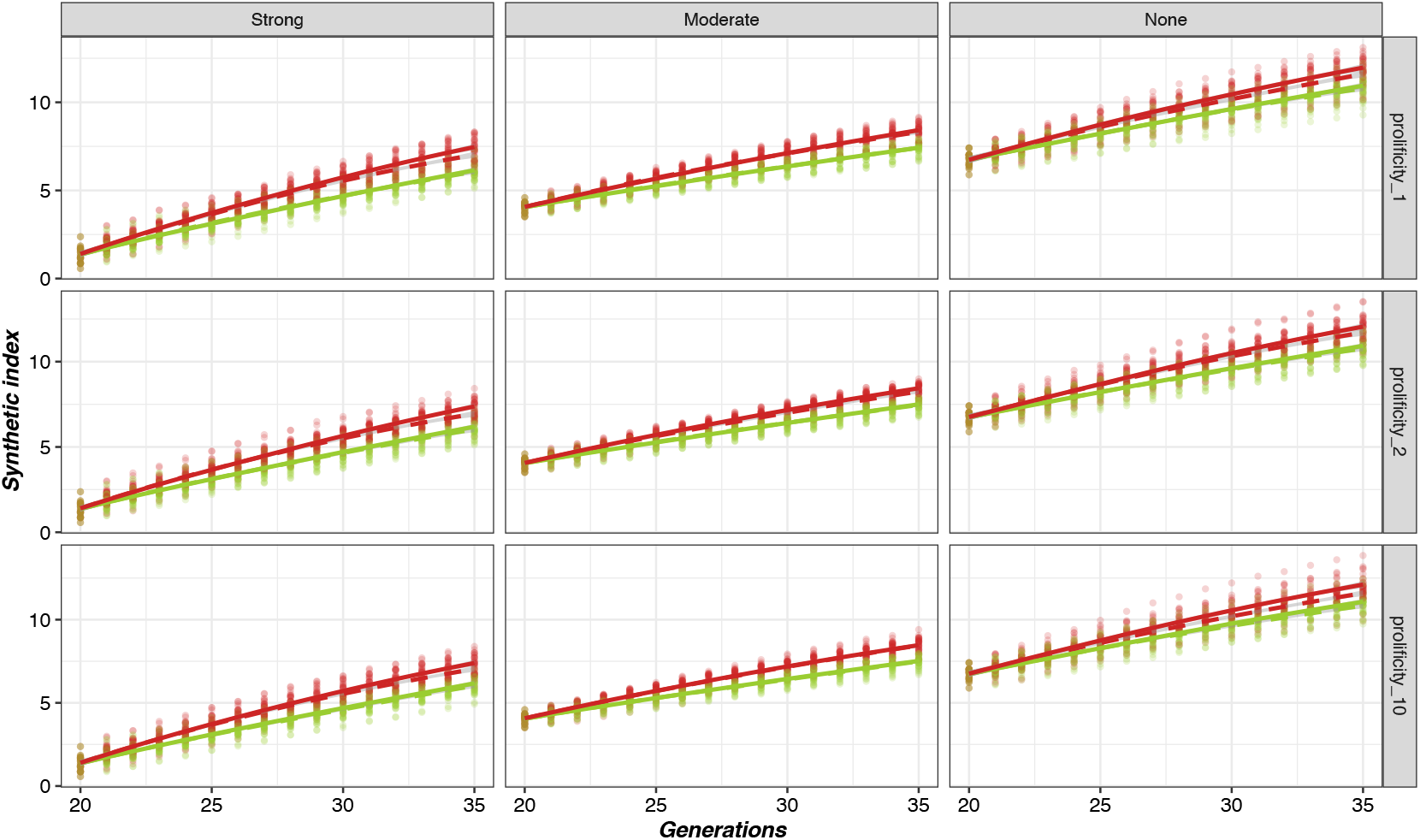
Evolution over time of the average estimated genetic value of the synthetic index of populations in the different scenarios, with and without the use of cryopreserved collections. In red, the pop_max_BV program, and in green, the pop_OCS program. Solid and dashed lines correspond respectively to scenarios without and with the use of *ex situ* collections.

**Figure 10.**
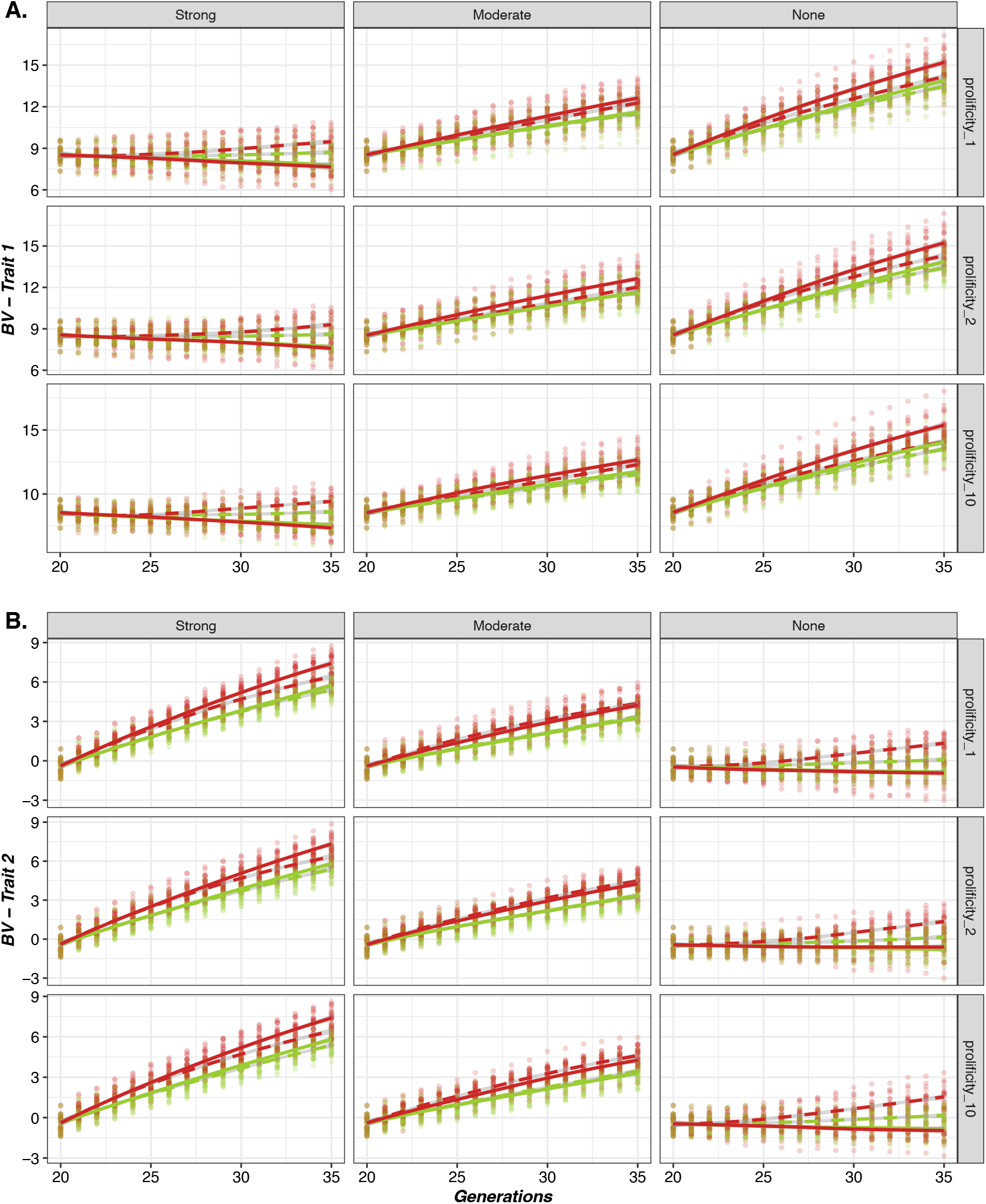
Evolution over time of the average genetic value of populations for the two traits of interest in the different scenarios, with and without the use of cryopreserved collections. In red, the pop_max_BV program, and in green, the pop_OCS program. Solid and dashed lines correspond respectively to scenarios without and with the use of *ex situ* collections.

For the pop_max_BV program, the use of *ex situ* resources had a positive, but weak, impact on the degradation of additive genetic variance, except when the selection objective was kept constant, in which case there was a more marked positive effect (see Figure 11). Prolificity had no significant effect on the loss of additive genetic variance for either of the two traits (three-factor ANOVA, F=1.11, df=2, p=0.33 for Trait 1 and F=1.20, df=2, p=0.30 for Trait 2). For the pop_OCS scenarios, the use of cryopreserved genetic resources had no significant impact on the additive genetic variance of either trait (F=0.10, df=1, p=0.76 for Trait 1 and F=0.60, df=1, p=0.44 for Trait 2).

**Figure 11.**
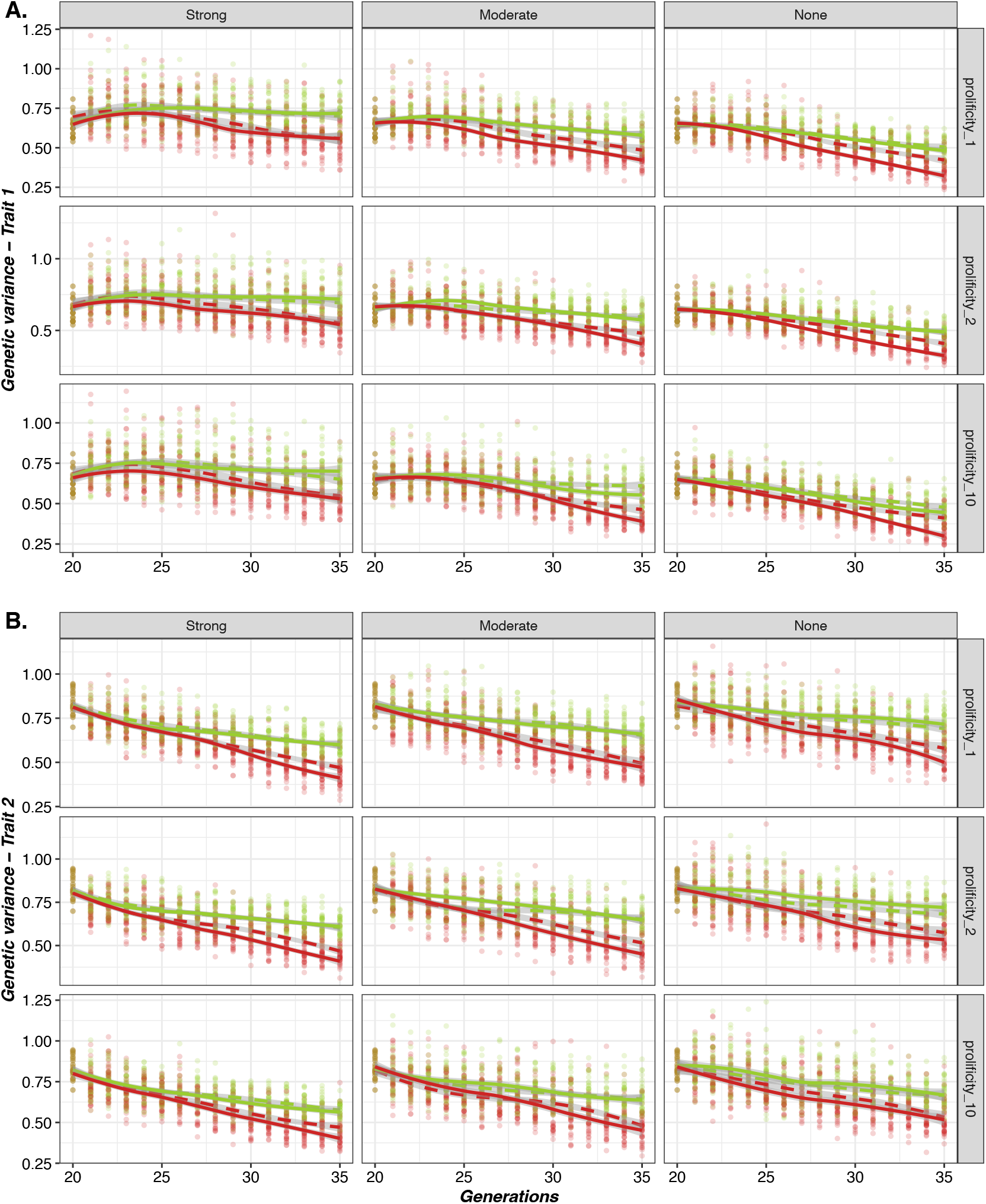
Evolution over time of the average additive genetic variance of populations for the two traits in the different scenarios, with and without the use of cryopreserved collections. In red, the pop_max_BV program, and in green, the pop_OCS program. Solid and dashed lines correspond respectively to scenarios without and with the use of *ex situ* collections.

## Discussion

Our study quantified how the use of contemporary and cryopreserved sires over several generations could affect the genetic diversity and genetic progress of populations being managed either for selection or conservation purposes. To our knowledge, this work is the first to use stochastic simulations to model and optimize the use of cryopreserved genetic material over multiple generations; previous research has focused on either deterministic simulations or real data analysis addressing the case of a single generation (Leroy et al., 2011; Doekes et al., 2018; Eynard et al., 2018).

We observed major differences between the scenarios depending on the breeding program employed (Table 2).

**Table 2.** Summary of the impact of the use of cryopreserved collections on breeding values and genetic variance for different breeding programs.

| PROGRAMS | Change in selection objective | Diversity | Breeding values |  |  | Genetic variance |  |
| --- | --- | --- | --- | --- | --- | --- | --- |
|  |  |  | Synthetic index | Trait 1 | Trait 2 | Trait 1 | Trait 2 |
| Random mating | NA | +++ | NA | NA | NA | NA | NA |
| Maximized genetic diversity | NA | +++ | NA | NA | NA | NA | NA |
| Optimal Contribution | None | ++ | = | - | ++ | = | = |
|  | Moderate | ++ | = | = | = | = | = |
|  | Strong | +++ | = | ++ | = | = | = |
| Maximized breeding value | None | + | = | -- | +++ | +++ | ++ |
|  | Moderate | = | = | - | = | ++ | ++ |
|  | Strong | = | = | +++ | - | + | ++ |

Note: NA=not applicable; '--' moderate deterioration; '-' small deterioration, '=' no change; '+' small improvement; '++' moderate improvement; '+++' large improvement

The results of our simulations confirm the effectiveness of using cryopreserved collections to maintain or reintroduce genetic diversity in animal populations. Even in the case of highly selected populations, the use of cryoconserved sires had a positive impact on genetic variance. Depending on the type of breeding program considered, then, the integration of cryopreserved materials makes it possible to maintain a high level of diversity or at least prevent its rapid deterioration.

This is especially obvious in conserved populations, where the main objective was to maintain genetic diversity, either by random selection of sires, including those from *ex situ* collections, or with a strategy designed to maximize genetic diversity in matings. In these cases, the genetic distances between populations 15 generations apart (from 20 to 35) was extremely small, showing a limited loss of genetic diversity over time (i.e., reduced genetic drift). Furthermore, the structure of the genetic diversity evolved within the populations, exhibiting an increase in the frequency of rare alleles (i.e., MAF<0.05). As a result, there was an overall increase in expected heterozygosity (He). Such a pattern would be expected to have a positive impact in terms of adaptation to new environments, which—in light of climate change—would be strongly desirable. In addition, levels of observed heterozygosity also benefited from the use of cryopreserved resources. Combined with reduced levels of kinship, this can have a positive impact on the fitness of conserved populations by limiting inbreeding depression (Chapman et al., 2009; Szulkin et al., 2010).

For populations under selection, where the only aim is to improve breeding values, the use of cryopreserved individuals also had a positive impact on genetic diversity. The advantage of using cryoconserved collections is that it increases the number of selection candidates in the breeding stock, and thus limits genetic drift. This can be seen in our results by the decrease in population differentiation over time (i.e., lower Nei distances between the start and end of the program), the increase in observed heterozygosity, and a higher frequency of rare alleles. The increase in genetic diversity through the use of *ex situ* collections also had a direct impact on selection: introducing this “neutral” genetic diversity resulted in increased genetic variance, demonstrating that even without the intention of reintroducing genetic diversity, cryopreserved resources are useful for enabling a selection response in the long term. While the synthetic index did not change based on the introduction of old material, the impact on a given trait strongly differed depending on the change in breeding weights for that trait. In the case of a reversal in the importance of a trait (negatively correlated to the other trait) in the selection index, the use of old material prevented a strong degradation of the trait. Conversely, if there was no change in objective, the benefit of cryopreserved resources lay in improving or maintaining the lower priority trait which, without the introduction of diversity, would deteriorate due to the negative correlation with the higher priority trait. This suggests that the real-world benefits of cryopreserved materials to breeding populations may be all the more important, as numerous traits are under selection and their importance varies with changes in the environment. In the context of the agro-ecological transition—aimed at adaptation to climate change and concomitant changes in breeding practices—it is highly likely that the number of traits considered will increase, and that their relative importance in selection objectives will vary (Dumont et al., 2013b; Wezel et al., 2020; Ducos et al., 2021). Thus, cryopreserved genetic resources will be particularly useful for optimizing the transition of highly selected populations into agro-ecological systems.

Populations for which the selection objective was integrated with a constraint on maintaining genetic diversity appeared to benefit less from the contribution of cryopreserved collections. There was no impact of their use on genetic variance, regardless of a change in selection objective. This result can be explained by the efficiency of the optimal contribution strategy in combining genetic improvement while maintaining genetic diversity. Regarding breeding values, though, the situation was different: the use of cryoconserved collections had a positive impact on the lower priority trait in the case of a constant selection objective, as well as on the higher priority trait in the case of a strong change in selection objective. Indeed, the use of OCS based on genomic data makes it possible to select the cryopreserved individuals that are most relevant for managing changes in breeding objectives, both in terms of maximizing genetic gain and by taking into account a constraint on increasing relatedness.

The number of males used from the cryobank varied among the breeeding programs. For populations under conservation, all individuals in the collection were used regularly during the program from generation 20 to generation 35. Both the OCS and random strategies yielded generally similar results, although the random strategy, strictly from the viewpoint of conservation of genetic diversity, would have a higher impact and might therefore be preferred (e.g., the random strategy avoided matings between closely related individuals). For highly selected populations, only a few individuals from collections were used, but it was possible for them to have large numbers of descendants in the subsequent generations, i.e., higher values than those obtained with OCS. This strategy ensures that a sufficiently large panel of offspring is available to be used as parents, preserving the re-introduced lineage over time. This utilization approach was reported in the reuse of an old sire within a cattle population under selection: frozen semen was used to produce several offspring, of which a few became confirmed sires in the breed (Jacques et al., 2023). Indeed, the larger the number of progeny produced by a single male, the more variability resulting from meiosis is obtained (i.e., gametic variance), thus increasing the chance of obtaining more favorable allelic combinations. This is especially likely when the donor exhibits a high level of heterozygosity.

In the present study, prolificity did not change the impact of using cryoconserved collections on either genetic diversity or breeding values. Because of this, our conclusions regarding the use of cryoconserved resources should remain valid across a range of livestock species with different litter sizes. However, prolificacy did have an impact on the number of cryopreserved sires used. This can be explained by the constant size of the simulated population, which sets a fixed number of offspring. For a given number of offspring, the number of artificial inseminations will differ based on litter size. This imposes a strong constraint on the female pathway when prolificity is high, leading to a reduction in the number of dams and matings and thus the possiblity of adjusting genetic contributions by OCS.

Some parameters that were not included in our simulations deserve further study. First, in the present study both simulated traits had the same heritability. It would be interesting to analyze a scenario in which heritability differs between traits, which is generally the case in different livestock species. Second, as mentioned above, it would be informative to specifically examine the impact of prolificity on the use of cryopreserved collections. For this, it would be necessary to compare breeding schemes with a fixed number of sires instead of a fixed number of offspring, as we chose to do here. Moreover, for our simulations we chose to use OCS, but other optimization methods are available. For example, another possibility for optimizing a breeding program is to use the minimum genomic parentage selection method (i.e., SPMG), which sets the desired genetic gain and aims to maximize genetic variability in the next generation (Colleau et al., 2004). This method is already being used in France for managing populations of goats under selection, such as Alpine and Saanen goats (Palhière et al., 2022). There are also similar methods for optimizing selection with diversity constraints in poultry (Chapuis et al., 2016). In the present study, we did not investigate the impact of using older material on the accuracy of genomic evaluations and therefore on selection efficiency. Indeed, for an accurate evaluation, the training (or reference) population must be similar to the population to be evaluated, as the estimation of marker effects relies on the link with causal alleles (Rincent et al., 2012); if the structure of the linkage disequilibrium changes, the effect of the markers may no longer be correctly predicted. As this structure evolves over time, it is likely that older individuals will be more disconnected from the current population, as well as from the reference population, which should remain contemporary. It would therefore be important to understand (i) the extent to which this could lead to a loss of accuracy in the assessments of older individuals and (ii) what kind of impact mixing individuals with more-or less-accurate predictions would have on population evolution. Another point to consider is that here we simulated a trait that can be measured directly on individuals, but a number of traits are measured indirectly. For phenotypes that are not directly measurable from breeding stock (e.g., a sire’s milk value estimated from the performance of his daughters, egg production in roosters), the accuracy of the estimation of genetic value depends mainly on measurements made on related individuals, and therefore on the number of offspring produced. In this case, the animals with the best-characterized phenotypes would be the oldest individuals, who would be expected to have the most accurate evaluations. These questions should be explored in greater detail with future simulations, so that new recommendations may be provided for the evaluation of cryopreserved genetic resources going forward.

Overall, our simulations demonstrate that, given the same cryobank architecture, the best use of cryopreserved individuals can be drastically different depending on the type of breeding scheme in place. Moreover, we did not consider in our simulations any constraints or limits on the number of available insemination doses, which may not be the case for all breeds or males (as demonstrated by an analysis of the French National Cryobank; Jacques et al., 2024). Conserved populations require management that optimizes genetic diversity, including the use of cryopreserved genetic resources at each generation, while populations under selection would be more likely to require periodic inputs from germplasm collections, in which the choice of individuals would focus predominantly on performance for a combination of traits that may change over time. In addition, here the use of cryopreserved sires in the various selection schemes was quite high across generations, which would require the availability of a sufficient number of doses (e.g., straws) in collections. If cryopreserved collections are used to meet different objectives, the ideal constitution of the collection, if it exists, would have to be defined according to the management program of each population. Further studies are needed to investigate the suitability of different strategies for developing germplasm collections. Indeed, although cryopreserved genetic resources can be stored indefinitely, the collection and storage of such materials generate costs that must be controlled (De Oliveira Silva et al., 2019). It is therefore necessary to develop strategies for managing the content of cryobanks so that (i) they remain relevant for a range of populations over time and (ii) managers of cryopreserved *ex situ* genetic resources retain some degree of freedom regarding the sampling and distribution of collections.

## Conclusions

In conclusion, the use of cryoconserved collections was shown here to be beneficial for the management of genetic diversity in the long term in all of the scenarios investigated. For populations under selection, the use of cryobank resources should be particularly useful in the case of a radical change in selection objectives. For this, the occasional use of a few sires, judiciously chosen so as not to affect genetic progress too much, should be sufficient. For populations under conservation, cryobanks bring major benefits, as the regular use of *ex situ* genetic resources could slow down the erosion of genetic variability. However, when germplasm collections are employed in these situations, it is important that the number of offspring generated is appropriate given the size of the population, which implies having a sufficient stock of material. Future studies should examine the value of cryobanks in a larger set of breeding schemes, particularly from the viewpoint of the agro-ecological transition, which could create new prospects for the better exploitation of cryopreserved genetic resources.

## Acknowledgments

We are grateful to the INRAE MIGALE bioinformatics facility (MIGALE, INRAE, 2020. Migale Bioinformatics Facility, doi: 10.15454/1.5572390655343293E12) for the help and computing resources provided. The authors would like to acknowledge Torsten Pook for his assistance with the MoBPS software.

## Authors’ contributions

GR and MTB conceived the project. AJ performed the analysis and drafted the manuscript. GR and MTB supervised the analyses. AJ, MTB, and GR interpreted the results. AJ, GR, and MTB contributed to the writing of the manuscript. All authors read and approved the final manuscript.

## References

Altieri, M. A., C. I. Nicholls, A. Henao, and M. A. Lana. 2015. Agroecology and the design of climate change-resilient farming systems. Agron. Sustain. Dev. 35:869–890. doi:10.1007/s13593-015-0285-2.

Chang, C. C., C. C. Chow, L. C. Tellier, S. Vattikuti, S. M. Purcell, and J. J. Lee. 2015. Second-generation PLINK: rising to the challenge of larger and richer datasets. GigaScience. 4:7. doi:10.1186/s13742-015-0047-8.

Chapman, J. R., S. Nakagawa, D. W. Coltman, J. Slate, and B. C. Sheldon. 2009. A quantitative review of heterozygosity–fitness correlations in animal populations. Molecular Ecology. 18:2746–2765. doi:10.1111/j.1365-294X.2009.04247.x.

Chapuis, H., C. Pincent, and J. j. Colleau. 2016. Optimizing selection with several constraints in poultry breeding. Journal of Animal Breeding and Genetics. 133:3–12. doi:10.1111/jbg.12178.

Colleau, J.-J., S. Moureaux, M. Briend, and J. Bechu. 2004. A method for the dynamic management of genetic variability in dairy cattle. Genetics Selection Evolution. 36:373. doi:10.1186/1297-9686-36-4-373.

Danchin Burge, C., E. Verrier, S. Moureaux, M. Tixier-Boichard, and B. Bibe. 2006. Sampling strategies and overview of the French national cryobank collections. In: 8th World Congress on Genetics Applied to Livestock Production.

Belo Horizonte, M., Brasil. De Oliveira Silva R., B. V. Ahmadi, S. J. Hiemstra, and D. Moran. 2019. Optimizing ex situ genetic resource collections for European livestock conservation. J Anim Breed Genet. 136:63–73. doi:10.1111/jbg.12368.

Doekes, H. P., R. F. Veerkamp, P. Bijma, S. J. Hiemstra, and J. Windig. 2018. Value of the Dutch Holstein Friesian germplasm collection to increase genetic variability and improve genetic merit. J Dairy Sci. 101:10022–10033. doi:10.3168/jds.2018-15217.

Doekes, H. P., R. F. Veerkamp, P. Bijma, G. de Jong, S. J. Hiemstra, and J. J. Windig. 2019. Inbreeding depression due to recent and ancient inbreeding in Dutch Holstein–Friesian dairy cattle. Genet Sel Evol. 51:54. doi:10.1186/s12711-019-0497-z.

Doublet, A.-C., P. Croiseau, S. Fritz, A. Michenet, C. Hozé, C. Danchin-Burge, D. Laloë, and G. Restoux. 2019. The impact of genomic selection on genetic diversity and genetic gain in three French dairy cattle breeds. Genet Sel Evol. 51:52. doi:10.1186/s12711-019-0495-1.

Ducos, A., F. Douhard, D. Savietto, M. Sautier, V. Fillon, M. Gunia, R. Rupp, C. Moreno-Romieux, S. Mignon-Grasteau, H. Gilbert, and L. Fortun-Lamothe. 2021. Contribution of animal genetics to the agroecological transition of livestock farming systems. INRAE Productions Animales. 34:79–96. doi:10.20870/productions-animales.2021.34.2.4773.

Dumont, B., L. Fortun-Lamothe, M. Jouven, M. Thomas, and M. Tichit. 2013a. Prospects from agroecology and industrial ecology for animal production in the 21st century. animal. 7:1028–1043. doi:10.1017/S1751731112002418.

Dumont, B., L. Fortun-Lamothe, M. Jouven, M. Thomas, and M. Tichit. 2013b. Prospects from agroecology and industrial ecology for animal production in the 21st century. Animal. 7:1028–1043. doi:10.1017/S1751731112002418.

Dumont, B., L. Puillet, G. Martin, D. Savietto, J. Aubin, S. Ingrand, V. Niderkorn, L. Steinmetz, and M. Thomas. 2020. Incorporating Diversity Into Animal Production Systems Can Increase Their Performance and Strengthen Their Resilience. Frontiers in Sustainable Food Systems. 4. Available from: https://www.frontiersin.org/articles/10.3389/fsufs.2020.00109

Escribano, A. J. 2016. Organic Livestock Farming — Challenges, Perspectives, and Strategies to Increase Its Contribution to the Agrifood System’s Sustainability — A Review. IntechOpen. Available from: https://www.intechopen.com/state.item.id

Eynard, S. E., J. J. Windig, S. J. Hiemstra, and M. P. L. Calus. 2016. Whole-genome sequence data uncover loss of genetic diversity due to selection. Genetics Selection Evolution. 48:33. doi:10.1186/s12711-016-0210-4.

Eynard, S. E., J. J. Windig, I. Hulsegge, S. J. Hiemstra, and M. P. L. Calus. 2018. The impact of using old germplasm on genetic merit and diversity – a cattle breed case study. J Anim Breed Genet. 135:311–322. doi:10.1111/jbg.12333.

FAO. 1998. Secondary Guidelines for Development of National Farm Animal Genetic Resources Management Plans – Management of small populations at risk. FAO, Rome, Italy. Available from: https://www.fao.org/documents/card/fr/c/d074fc39-aca0-58ab-8024-beaf80ba6014/

Fox, J., S. Weisberg, B. Price, D. Adler, D. Bates, G. Baud-Bovy, B. Bolker, S. Ellison, D. Firth, M. Friendly, G. Gorjanc, S. Graves, R. Heiberger, P. Krivitsky, R. Laboissiere, M. Maechler, G. Monette, D. Murdoch, H. Nilsson, D. Ogle, B. Ripley, T. Short, W. Venables, S. Walker, D. Winsemius, A. Zeileis, and R-Core. 2019. car: Companion to Applied Regression. https://cran.r-project.org/web/packages/car/index.html. Accessed 24 Oct 2022.

Gaughan, J. B., V. Sejian, T. L. Mader, and F. R. Dunshea. 2019. Adaptation strategies: ruminants. Animal Frontiers. 9:47–53. doi:10.1093/af/vfy029.

Hagger, C. 2005. Estimates of genetic diversity in the brown cattle population of Switzerland obtained from pedigree information. Journal of Animal Breeding and Genetics. 122:405–413. doi:10.1111/j.1439-0388.2005.00552.x.

Hoffmann, I. 2010. Climate change and the characterization, breeding and conservation of animal genetic resources. Animal Genetics. 41:32–46. doi:10.1111/j.1365-2052.2010.02043.x.

Hulsegge, I., K. Oldenbroek, A. Bouwman, R. Veerkamp, and J. Windig. 2022. Selection and Drift: A Comparison between Historic and Recent Dutch Friesian Cattle and Recent Holstein Friesian Using WGS Data. Animals. 12:329. doi:10.3390/ani12030329.

IPES-Food. 2016. From uniformity to diversity: A paradigm shift from industrial agriculture to diversified agroecological systems. IPES-Food (Independent Panel of Experts on Sustainable Food Systems).

Jacques, A., D. Duclos, C. Danchin-Burge, M.-J. Mercat, M. Tixier-Boichard, and G. Restoux. 2024. Assessing the potential of germplasm collections for the management of genetic diversity: the case of the French National Cryobank. Peer Community Journal. 4:e13. doi:10.24072/pcjournal.369.

Jacques, A., G. Leroy, X. Rognon, E. Verrier, M. Tixier-Boichard, and G. Restoux. 2023. Reintroducing genetic diversity in populations from cryopreserved material: the case of Abondance, a French local dairy cattle breed. Genet Sel Evol. 55:28. doi:10.1186/s12711-023-00801-6.

Kantanen, J., P. Løvendahl, E. Strandberg, E. Eythorsdottir, M.-H. Li, A. Kettunen-Præbel, P. Berg, and T. Meuwissen. 2015. Utilization of farm animal genetic resources in a changing agro-ecological environment in the Nordic countries. Frontiers in Genetics. 6. Available from: https://www.frontiersin.org/articles/10.3389/fgene.2015.00052

Lenth, R. V., P. Buerkner, M. Herve, M. Jung, J. Love, F. Miguez, H. Riebl, and H. Singmann. 2021. emmeans: Estimated Marginal Means, aka least-squares means. https://CRAN.R-project.org/package=emmeans/. Accessed 24 Oct 2022.

Leroy, G., C. Danchin-Burge, and E. Verrier. 2011. Impact of the use of cryobank samples in a selected cattle breed: a simulation study. Genet Sel Evol. 43:36. doi:10.1186/1297-9686-43-36.

Meuwissen, T. H. 1997. Maximizing the response of selection with a predefined rate of inbreeding. J Anim Sci. 75:934–40. doi:10.2527/1997.754934x.

Nei, M. 1972. Genetic Distance between Populations. The American Naturalist. 106:283–292. doi:10.1086/282771.

Notter, D. R. 1999. The importance of genetic diversity in livestock populations of the future. J Anim Sci. 77:61–69. doi:10.2527/1999.77161x.

Palhière, I., V. Gousseau, and J.-J. Colleau. 2022. La Sélection à Parenté Minimum Génomique: principes et résultats pour les deux races caprines laitières principales françaises. INRAE Productions Animales. 35:171–186. doi:10.20870/productions-animales.2022.35.3.7152.

Pasqui, M., and E. Di Giuseppe. 2019. Climate change, future warming, and adaptation in Europe. Anim Front. 9:6–11. doi:10.1093/af/vfy036.

Pook, T. 2021. MoBPS: Modular Breeding Program Simulator. Available from: https://cran.r-project.org/web/packages/MoBPS/index.html

Pook, T., M. Schlather, and H. Simianer. 2020. MoBPS – Modular Breeding Program Simulator. G3: Genes, Genomes, Genetics. 10:1915–1918. doi:10.1534/g3.120.401193.

Purcell, S., B. Neale, K. Todd-Brown, L. Thomas, M. A. R. Ferreira, D. Bender, J. Maller, P. Sklar, P. I. W. de Bakker, M. J. Daly, and P. C. Sham. 2007. PLINK: A Tool Set for Whole-Genome Association and Population-Based Linkage Analyses. The American Journal of Human Genetics. 81:559–575. doi:10.1086/519795.

R Core team. 2020. R: a language and environment for statistical computing.

Rincent, R., D. Laloë, S. Nicolas, T. Altmann, D. Brunel, P. Revilla, V. M. Rodríguez, J. Moreno-Gonzalez, A. Melchinger, E. Bauer, C.-C. Schoen, N. Meyer, C. Giauffret, C. Bauland, P. Jamin, J. Laborde, H. Monod, P. Flament, A. Charcosset, and L. Moreau. 2012. Maximizing the Reliability of Genomic Selection by Optimizing the Calibration Set of Reference Individuals: Comparison of Methods in Two Diverse Groups of Maize Inbreds (Zea mays L.). Genetics. 192:715–728. doi:10.1534/genetics.112.141473.

Sonesson, A. K., J. A. Woolliams, and T. H. Meuwissen. 2012. Genomic selection requires genomic control of inbreeding. Genet Sel Evol. 44:27. doi:10.1186/1297-9686-44-27.

Szulkin, M., N. Bierne, and P. David. 2010. Heterozygosity-fitness correlations: a time for reappraisal. Evolution. doi:10.1111/j.1558-5646.2010.00966.x. Available from: https://academic.oup.com/evolut/article/64/5/1202/6854403

Thornton, P. K. 2010. Livestock production: recent trends, future prospects. Philosophical Transactions of the Royal Society B: Biological Sciences. 365:2853–2867. doi:10.1098/rstb.2010.0134.

VanRaden, P. M. 2008. Efficient Methods to Compute Genomic Predictions. Journal of Dairy Science. 91:4414–4423. doi:10.3168/jds.2007-0980.

Wellmann, R. 2019. Optimum contribution selection for animal breeding and conservation: the R package optiSel. BMC Bioinformatics. 20:25. doi:10.1186/s12859-018-2450-5.

Wellmann, R. 2021. optiSel: Optimum Contribution Selection and Population Genetics. Available from: https://CRAN.R-project.org/package=optiSel

Wezel, A., B. G. Herren, R. B. Kerr, E. Barrios, A. L. R. Gonçalves, and F. Sinclair. 2020. Agroecological principles and elements and their implications for transitioning to sustainable food systems. A review. Agron. Sustain. Dev. 40:40. doi:10.1007/s13593-020-00646-z.

Wickham, H. 2011. ggplot2. Wiley Interdiscip Rev Comput Stat. 3:180–185. doi:10.1002/wics.147.

