## Supplementary figures and images for "Optimizing the use of cryopreserved genetic resources for the selection and conservation of animal populations"

### Additional File 1 Figure S1

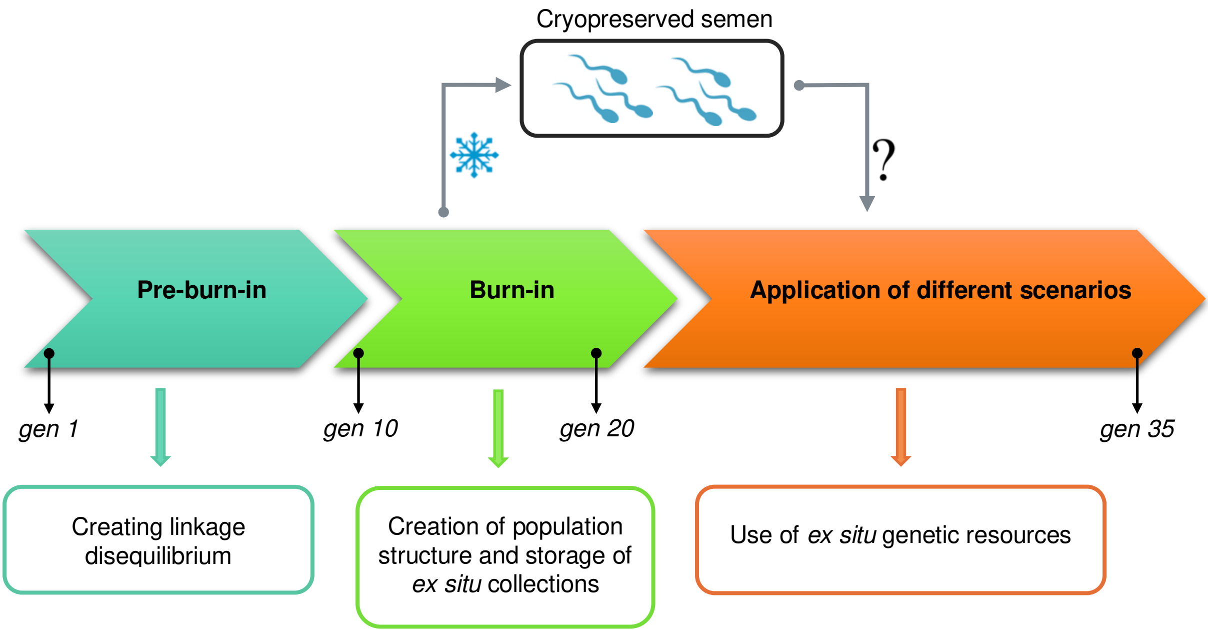
