## Additional File 2 for "Optimizing the use of cryopreserved genetic resources for the selection and conservation of animal populations"

**Table S1:** Measures of genetic diversity evolution for pop_max_BV scenarios.

| SCENARIO TYPE | Without use of cryopreserved genetic resources | | | | | With use of cryopreserved genetic resources | | | | |
|---|---|---|---|---|---|---|---|---|---|---|
| | **ΔKinship** | **ΔHet** | **ΔFreq** | **Genetic distance** | **No. enriched rare SNP** | **ΔKinship** | **ΔHet** | **ΔFreq** | **Genetic distance** | **No. enriched rare SNP** |
| **Prolificity 1** | | | | | | | | | | |
| None | 2.78E-01 | -9.83E-02 | -7.86E-02 | 6.62E-02 | 51 | 1.98E-01 | -6.96E-02 | -5.38E-02 | 4.55E-02 | 66 |
| *sd* | *2.69E-02* | *1.08E-02* | *1.01E-02* | *8.00E-03* | *7* | *2.27E-02* | *8.29E-03* | *6.77E-03* | *5.74E-03* | *10* |
| Moderate | 2.54E-01 | -8.89E-02 | -7.14E-02 | 6.22E-02 | 57 | 2.14E-01 | -7.45E-02 | -5.84E-02 | 5.15E-02 | 67 |
| *sd* | *2.28E-02* | *8.25E-03* | *7.97E-03* | *6.50E-03* | *9* | *2.04E-02* | *7.62E-03* | *6.69E-03* | *5.14E-03* | *9* |
| Strong | 2.64E-01 | -9.17E-02 | -7.41E-02 | 7.10E-02 | 61 | 2.06E-01 | -7.25E-02 | -5.62E-02 | 5.25E-02 | 70 |
| *sd* | *2.96E-02* | *1.12E-02* | *1.03E-02* | *8.80E-03* | *9* | *2.26E-02* | *8.24E-03* | *7.22E-03* | *4.75E-03* | *12* |
| **Prolificity 2** | | | | | | | | | | |
| None | 2.80E-01 | -9.74E-02 | -7.91E-02 | 6.88E-02 | 55 | 2.06E-01 | -7.14E-02 | -5.57E-02 | 4.77E-02 | 64 |
| *sd* | *2.64E-02* | *1.02E-02* | *1.04E-02* | *7.37E-03* | *10* | *1.93E-02* | *6.77E-03* | *6.21E-03* | *6.96E-03* | *8* |
| Moderate | 2.66E-01 | -9.22E-02 | -7.42E-02 | 6.59E-02 | 55 | 2.07E-01 | -7.18E-02 | -5.61E-02 | 4.98E-02 | 65 |
| *sd* | *1.98E-02* | *7.44E-03* | *6.60E-03* | *6.68E-03* | *8* | *1.92E-02* | *7.32E-03* | *6.89E-03* | *4.61E-03* | *5* |
| Strong | 2.56E-01 | -8.90E-02 | -7.20E-02 | 6.86E-02 | 60 | 2.00E-01 | -6.90E-02 | -5.32E-02 | 5.13E-02 | 71 |
| *sd* | *2.88E-02* | *1.02E-02* | *1.08E-02* | *1.04E-02* | *8* | *1.94E-02* | *7.39E-03* | *5.95E-03* | *4.48E-03* | *10* |
| **Prolificity 10** | | | | | | | | | | |
| None | 2.95E-01 | -1.02E-01 | -8.16E-02 | 7.08E-02 | 47 | 2.24E-01 | -7.75E-02 | -6.17E-02 | 5.19E-02 | 63 |
| *sd* | *2.42E-02* | *1.01E-02* | *8.56E-03* | *8.25E-03* | *7* | *2.45E-02* | *9.06E-03* | *8.10E-03* | *5.31E-03* | *9* |
| Moderate | 2.81E-01 | -9.77E-02 | -7.88E-02 | 6.83E-02 | 51 | 2.24E-01 | -7.79E-02 | -6.07E-02 | 5.43E-02 | 60 |
| *sd* | *3.11E-02* | *1.06E-02* | *9.96E-03* | *8.56E-03* | *7* | *2.13E-02* | *7.58E-03* | *5.81E-03* | *5.35E-03* | *8* |
| Strong | 2.69E-01 | -9.38E-02 | -7.67E-02 | 7.23E-02 | 60 | 2.27E-01 | -7.84E-02 | -6.22E-02 | 577E-02 | 68 |
| *sd* | *2.79E-02* | *1.06E-02* | *9.73E-03* | *8.00E-03* | *10* | *3.12E-02* | *1.19E-02* | *1.05E-02* | *7.54E-03* | *11* |

Note: All abbreviations correspond to those developed in the Method section.

None: scenario with no change in the weighting of the two traits - Moderate: weighting of both traits increased to 0.5 - Strong: complete inversion of weights between the two traits

ΔKinship: the delta between kinship in generations 20 and 35 - ΔHet: the delta between heterozygosity in generations 20 and 35

ΔFreq: the delta between the allelic frequencies of the generation 20 and generation 35

No. enriched rare SNP corresponds to the number of SNPs with a MAF lower than 0.05 in generation 20 and whose frequency increased in generation 35.

**Table S2:** Measures of genetic diversity evolution for pop_OCS scenarios.

| SCENARIO TYPE | Without use of cryopreserved genetic resources | | | | | With use of cryopreserved genetic resources | | | | |
|---|---|---|---|---|---|---|---|---|---|---|
| | **ΔKinship** | **ΔHet** | **ΔFreq** | **Genetic distance** | **No. enriched rare SNP** | **ΔKinship** | **ΔHet** | **ΔFreq** | **Genetic distance** | **No. enriched rare SNP** |
| **Prolificity 1** | | | | | | | | | | |
| None | 9.28E-02 | -3.43E-02 | -2.52E-02 | 1.89E-02 | 107 | 7.50E-02 | -2.72E-02 | -1.99E-02 | 1.38E-02 | 116 |
| *sd* | *1.12E-02* | *4.29E-03* | *3.99E-03* | *2.47E-03* | *8* | *7.05E-03* | *2.85E-03* | *2.27E-03* | *9.51E-04* | *15* |
| Moderate | 8.27E-02 | -3.00E-02 | -2.26E-02 | 1.82E-02 | 116 | 7.59E-02 | -2.76E-02 | -2.03E-02 | 1.59E-02 | 123 |
| *sd* | *9.74E-03* | *3.97E-03* | *2.90E-03* | *1.56E-03* | *13* | *6.95E-03* | *2.72E-03* | *2.43E-03* | *1.43E-03* | *13* |
| Strong | 7.42E-02 | -2.70E-02 | -2.02E-02 | 1.99E-02 | 127 | 6.88E-02 | -2.55E-02 | -1.85E-02 | 1.74E-02 | 132 |
| *sd* | *7.16E-03* | *3.60E-03* | *3.25E-03* | *1.65E-03* | *11* | *8.28E-03* | *2.89E-03* | *2.57E-03* | *1.62E-03* | *14* |
| **Prolificity 2** | | | | | | | | | | |
| None | 9.45E-02 | -3.41E-02 | -2.55E-02 | 1.92E-02 | 108 | 8.08E-02 | -2.94E-02 | -2.16E-02 | 1.49E-02 | 110 |
| *sd* | *8.39E-03* | *3.34E-03* | *2.89E-03* | *2.02E-03* | *11* | *6.67E-03* | *2.60E-03* | *2.07E-03* | *9.97E-04* | *7* |
| Moderate | 8.79E-02 | -3.09E-02 | -2.30E-02 | 1.94E-02 | 115 | 8.15E-02 | -2.95E-02 | -2.16E-02 | 1.73E-02 | 114 |
| *sd* | *1.04E-02* | *4.91E-03* | *4.34E-03* | *2.48E-03* | *13* | *8.81E-03* | *3.92E-03* | *3.68E-03* | *1.52E-03* | *10* |
| Strong | 8.15E-02 | -2.96E-02 | -2.18E-02 | 2.16E-02 | 124 | 7.00E-02 | -2.53E-02 | -1.83E-02 | 1.80E-02 | 128 |
| *sd* | *9.32E-03* | *3.70E-03* | *3.25E-03* | *2.36E-03* | *14* | *7.10E-03* | *3.12E-03* | *2.27E-03* | *1.43E-03* | *16* |
| **Prolificity 10** | | | | | | | | | | |
| None | 1.36E-01 | -4.87E-02 | -3.68E-02 | 2.95E-02 | 88 | 1.03E-01 | -3.69E-02 | -2.77E-02 | 2.10E-02 | 103 |
| *sd* | *1.42E-02* | *5.45E-03* | *4.47E-03* | *3.02E-03* | *12* | *1.06E-02* | *4.61E-03* | *4.50E-03* | *2.43E-03* | *13* |
| Moderate | 1.21E-01 | -4.24E-02 | -3.21E-02 | 2.74E-02 | 97 | 1.07E-01 | -3.83E-02 | -2.85E-02 | 2.38E-02 | 101 |
| *sd* | *1.48E-02* | *5.36E-03* | *4.10E-03* | *3.32E-03* | *10* | *1.28E-02* | *5.05E-03* | *4.23E-03* | *2.80E-03* | *15* |
| Strong | 1.12E-01 | -3.93E-02 | -2.95E-02 | 3.00E-02 | 105 | 9.67E-02 | -3.45E-02 | -2.54E-02 | 2.46E-02 | 115 |
| *sd* | *1.28E-02* | *5.51E-03* | *4.66E-03* | *2.87E-03* | *10* | *1.77E-02* | *6.59E-03* | *5.49E-03* | *3.20E-03* | *14* |

Note: All abbreviations correspond to those developed in the Method section.
None: scenario with no change in the weighting of the two traits - Moderate: weighting of both traits increased to 0.5 - Strong: complete inversion of weights between the two traits
ΔKinship: the delta between kinship in generations 20 and 35 - ΔHet: the delta between heterozygosity in generations 20 and 35
ΔFreq: the delta between the allelic frequencies of the generation 20 and generation 35
No. enriched rare SNP corresponds to the number of SNPs with a MAF lower than 0.05 in generation 20 and whose frequency increased in generation 35.

**Table S3:** Measures of genetic diversity evolution for conservation scenarios.

| SCENARIO TYPE | | ΔKinship | ΔHet | ΔFreq | Genetic distance | No. enriched rare SNP |
|---|---|---|---|---|---|---|
| | | | | **Scenario pop_rm** | | |
| **Prolificity 1** | | | | | | |
| | no_cryo | 1,62E-02 | -5,69E-03 | -4,02E-03 | 4,23E-03 | 169 |
| | *sd* | *2,38E-03* | *1,27E-03* | *9,34E-04* | *2,32E-04* | *18* |
| | cryo | 5,37E-03 | -1,78E-03 | -1,11E-03 | 3,70E-03 | 231 |
| | *sd* | *2,18E-03* | *1,19E-03* | *1,18E-03* | *1,59E-04* | *22* |
| **Prolificity 2** | | | | | | |
| | no_cryo | 1,72E-02 | -5,85E-03 | -4,21E-03 | 4,56E-03 | 175 |
| | *sd* | *1,92E-03* | *1,26E-03* | *1,06E-03* | *2,82E-04* | *22* |
| | cryo | 5,72E-03 | -2,13E-03 | -1,42E-03 | 3,85E-03 | 230 |
| | *sd* | *2,73E-03* | *1,40E-03* | *1,06E-03* | *2,69E-04* | *25* |
| **Prolificity 10** | | | | | | |
| | no_cryo | 2,99E-02 | -9,66E-03 | -7,07E-03 | 7,72E-03 | 150 |
| | *sd* | *2,35E-03* | *1,52E-03* | *9,35E-04* | *4,75E-04* | *19* |
| | cryo | 1,02E-02 | -2,97E-03 | -2,45E-03 | 5,08E-03 | 217 |
| | *sd* | *3,36E-03* | *1,69E-03* | *1,02E-03* | *2,93E-04* | *22* |
| | | | | **Scenario pop_maxDG** | | |
| **Prolificity 1** | | | | | | |
| | no_cryo | 3,15E-02 | -1,18E-02 | -8,60E-03 | 7,01E-03 | 149 |
| | *sd* | *3,29E-03* | *1,88E-03* | *1,53E-03* | *3,24E-04* | *10* |
| | cryo | 7,62E-03 | -3,27E-03 | -2,01E-03 | 3,17E-03 | 230 |
| | *sd* | *2,27E-03* | *1,17E-03* | *8,19E-04* | *1,70E-04* | *19* |
| **Prolificity 2** | | | | | | |
| | no_cryo | 3,47E-02 | -1,28E-02 | -9,34E-03 | 7,76E-03 | 144 |
| | *sd* | *3,22E-03* | *2,26E-03* | *1,83E-03* | *5,19E-04* | *13* |
| | cryo | 8,57E-03 | -3,70E-03 | -2,48E-03 | 3,42E-03 | 226 |
| | *sd* | *2,70E-03* | *1,32E-03* | *9,47E-04* | *1,37E-04* | *22* |
| **Prolificity 10** | | | | | | |
| | no_cryo | 5,40E-02 | -1,86E-02 | -1,41E-02 | 1,23E-02 | 123 |
| | *sd* | *4,82E-03* | *2,36E-03* | *2,01E-03* | *9,42E-04* | *18* |
| | cryo | 1,80E-02 | -5,27E-03 | -4,51E-03 | 5,55E-03 | 199 |
| | *sd* | *2,88E-03* | *1,83E-03* | *1,47E-03* | *4,01E-04* | *20* |

Note: All abbreviations correspond to those developed in the Method section.
None: scenario with no change in the weighting of the two traits - Moderate: weighting of both traits increased to 0.5 - Strong: complete inversion of weights between the two traits
ΔKinship: the delta between kinship in generations 20 and 35 - ΔHet: the delta between heterozygosity in generations 20 and 35
ΔFreq: the delta between the allelic frequencies of the generation 20 and generation 35
No. enriched rare SNP corresponds to the number of SNPs with a MAF lower than 0.05 in generation 20 and whose frequency increased in generation 35.
